# A Transient Increase in Cardiomyocyte Protein O-GlcNAcylation Enhances Susceptibility to Pressure Overload-Induced Cardiac Remodeling

**DOI:** 10.1101/2025.06.04.657956

**Authors:** Samuel F. Chang, Chae-Myeong Ha, Manoja K. Brahma, Luke A. Potter, Mahima S. Reddy, Sayan Bakshi, Kerstin Preuß, Md Saimoon Rahman, Johannes K. Fischer, Caitlin A. Harrell, Zhihuan Sun, John C. Chatham, Adam R. Wende

**Affiliations:** Department of Pathology, Division of Molecular and Cellular Pathology, University of Alabama at Birmingham, Birmingham, AL, USA; Medical Scientist Training Program, Heersink School of Medicine, University of Alabama at Birmingham, Birmingham, AL, USA; Department of Cardiothoracic Surgery, University Hospital Jena, Friedrich-Schiller University of Jena, Jena, Germany

**Keywords:** cardiac remodeling, diabetic heart disease, metabolic memory, protein O-GlcNAcylation, pressure overload

## Abstract

**BACKGROUND:** The observation that diabetic patients always under tight-glycemic control consistently show better cardiovascular disease outcomes compared to patients who transition to tight-glycemic control after prior conventional glycemic control lead to the concept of metabolic memory. Mechanisms such as epigenetics possibly mediate the lasting metabolic memory effects, our understanding of the underlying mechanisms remains limited. Increased cardiac protein posttranslational O-linked β-N-acetylglucosamine (O-GlcNAc) modification is implicated in cardiac remodeling observed in diabetes, and our previous work shows chronically elevated cardiomyocyte O-GlcNAc causes adverse cardiac changes. Therefore, the current study hypothesized that transiently increased cardiomyocyte O-GlcNAcylation leads to exacerbated adverse cardiac remodeling after subsequent pressure-overload.

**METHODS AND RESULTS:** Using our previously described inducible cardiomyocyte specific, dominant-negative O-GlcNAcase (dnOGAh) mouse and single transgenic littermate controls (Con), we induced O-GlcNAc levels for 2wk (ON), followed by a 2wk washout (OFF); mice then underwent transverse-aortic constriction (TAC) or Sham surgery. We observed the expected cardiac remodeling in TAC groups, including decreased cardiac function, and increased hypertrophy and fibrosis. Moreover, these pathologic measures were exacerbated in the ON/OFF-TAC vs. Con-TAC mice; additionally, transcriptomic analysis of LV-tissue from each experimental group showed pathways which not only supported our fibrosis, hypertrophy and functional results of exacerbated cardiac remodeling, but also, revealed potential novel molecular pathways underlying this pathologic remodeling.

**CONCLUSIONS:** We observed exacerbated cardiac pathology between ON/OFF-TAC vs. Con-TAC groups supporting the concept of “O-GlcNAc memory” as a component of metabolic memory. Moreover, transcriptomic analysis provides insight into potential molecular pathways underpinning this metabolic/O-GlcNAc memory such as *Ccn2*/CTGF-driven fibrosis, and/or *Nox4*-driven oxidative stress.

**GRAPHICAL ABSTRACT:** 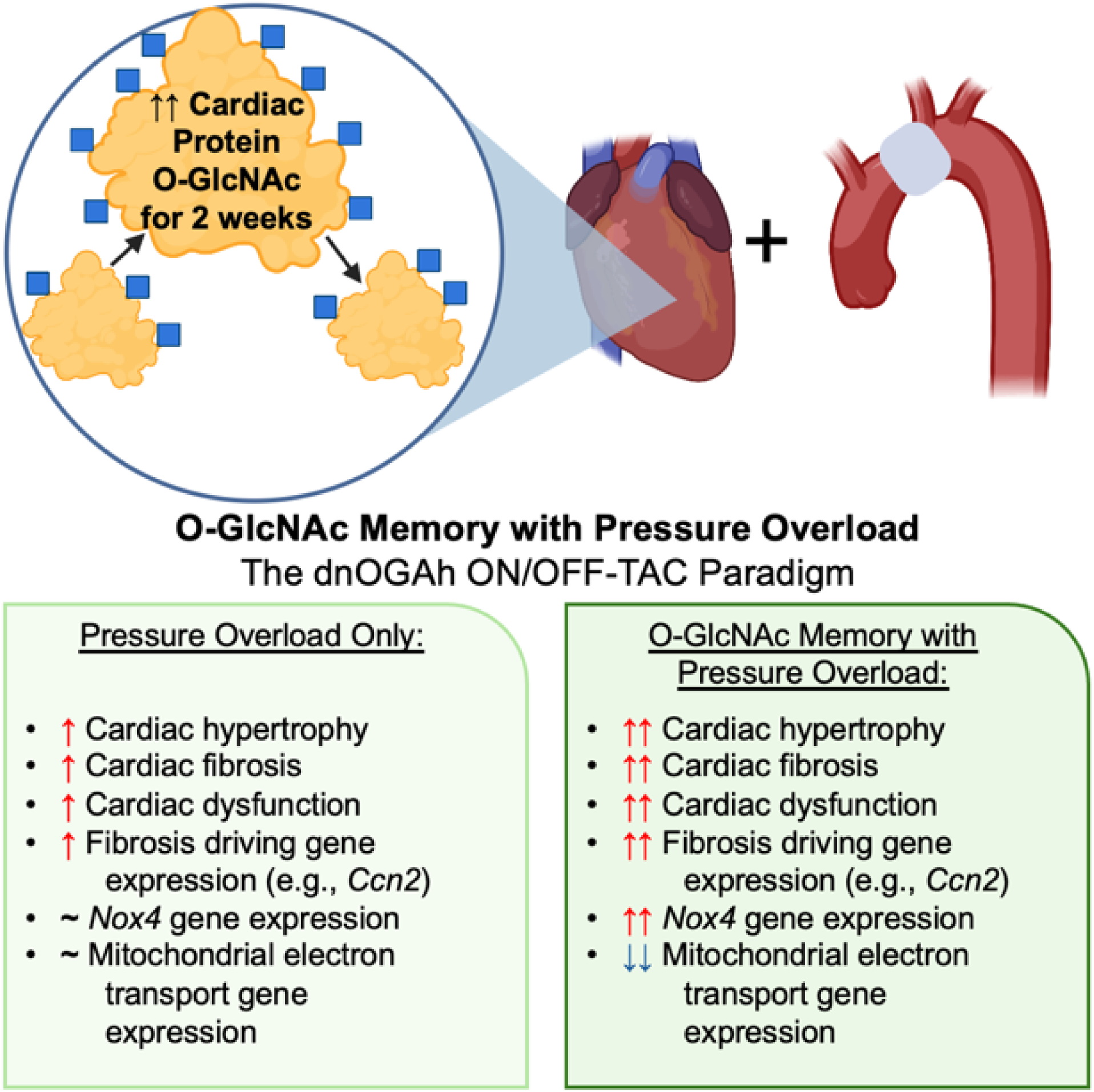

**Clinical Perspective:** *What is new?:*

- We provide a novel paradigm to study phenotypic and molecular effects of specific, transiently increased cardiomyocyte O-GlcNAcylation on the heart.
- Our results show exacerbated adverse cardiac remodeling due to transiently increased cardiomyocyte O-GlcNAc with pressure-overload, supporting the concept of “O-GlcNAc memory” as a component of metabolic memory.
- Transcriptomic insights show gene expression basis for not only observed exacerbated adverse cardiac remodeling (e.g., hypertrophy, fibrosis, cardiac dysfunction), but also potential molecular pathways that could drive cardiac pathology exacerbation of O-GlcNAc memory.

*What are the clinical implications?:*

- This study supports a concept of “O-GlcNAc memory”, where previously increased cardiomyocyte protein O-GlcNAcylation can impact the later development of differential cardiac pathology—like the pathology seen in metabolic memory research.
- The potential role of O-GlcNAc in mediating metabolic memory will help focus future translational research on this modification and downstream cardiac effects in diabetes.
- Transcriptomic profiling of cardiac remodeling in this model provides an investigational roadmap for future molecular and functional studies to identify novel therapeutics that ameliorate heart disease induced by differential metabolic memory.

## INTRODUCTION

Heart disease is the leading cause of death in the United States^1^ and globally.^2^ Additionally, prevalence of diabetes mellitus (diabetes/DM) is near 10% worldwide,^3^ and even higher in the United States;^4^ moreover, among the many health complications, cardiovascular diseases (CVD) are the number one cause of death in individuals with diabetes.^5^ Advancements in treatments have improved mortality outcomes, however, individuals with diabetes still have significantly increased CVD mortality even after accounting for other risk factors.^6-8^ The Diabetes Control and Complications Trial (DCCT) and later Epidemiology of Diabetes Interventions and Complications (EDIC) study found that patients always under tight glycemic control had significantly less major adverse cardiovascular events (MACE) compared to those initially on conventional therapy (during DCCT) then switched to tight glycemic control (EDIC study);^9-12^ this significant benefit has persisted at least 30 years.^13^ Results from DCCT/EDIC, along with studies such as the United Kingdom Prospective Diabetes Study (UKPDS) and Veterans Affair Diabetes Trial for type-2 diabetes, support the concept that previously different levels of glycemic control in diabetic patients could account for the subsequent long-term differences in cardiovascular outcomes–termed “metabolic memory”.^14-17^ Furthermore, studies have identified epigenetics as possible mechanisms behind metabolic memory’s differential outcomes.^18,19^

It is well known that the diabetes leads to severe perturbations of cardiac metabolism including cellular dysfunction in the context of lipotoxicity, glucotoxicity, and glucolipotoxicity;^20-27^ however, the exact biological mechanisms linking dysregulation of cardiac metabolism and cardiac dysfunction in the diabetes are not well understood. In both diabetic patient hearts and animal models, studies have shown protein post translational modifications (PTM) by O-linked β-N-acetylglucosamine (O-GlcNAc).^27,28^ O-GlcNAcylation occurs when O-GlcNAc transferase (OGT) adds a single O-GlcNAc to specific serine/threonine residues;^29^ removal is mediated by O-GlcNAcase (OGA).^30^ O-GlcNAc plays a variety of roles in physiology and pathology development;^31-33^ specifically in the heart, increased protein O-GlcNAc post-ischemia exerts a protective effect,^34-36^ while persistently increased cardiac protein O-GlcNAc can be cardiac maladaptive.^28,37^ Furthermore, in our recent study using a transgenic mouse model of increased cardiomyocyte protein O-GlcNAc to evaluate the specific effects of sustained cardiomyocyte O-GlcNAc on the heart, we found that chronically increased cardiomyocyte protein O-GlcNAc alone was sufficient to lead to cardiac pathology such as diastolic dysfunction, cardiac hypertrophy, fibrosis, and reduced mitochondrial function^38^, similar to the pathology seen in response to diabetes.

O-GlcNAc is also known to regulate both transcriptional and epigenetic regulation, and contribute to cellular memory;^31,39,40^ for example, O-GlcNAcylation of transcription factor Sp1 protects it from degradation^41,42^ and increases its transcriptional activity,^43^ and O-GlcNAcylation of ten-eleven translocation methyl-cytosine dioxygenases (TETs), responsible for active epigenetic DNA hydroxymethylation^44,45^, can lead to increased DNA hypomethylation.^46-48^ Collectively these provide the foundation to investigate whether “O-GlcNAc memory” may play a role in cardiac pathology and contribute to “metabolic memory”.

Thus, in our current study, we used the dnOGAh model to transiently increase cardiomyocyte protein O-GlcNAc levels for 2 weeks followed by a washout period and subsequently induce left ventricle pressure overload via transverse aortic constriction (TAC). We hypothesized that the transient increase in O-GlcNAc would exacerbate adverse cardiac remodeling under subsequent pressure-overload, consistent with the concept of O-GlcNAc memory. In functional, gravimetric, histologic and transcriptomic measures, we found overall exacerbated cardiac pathology in hearts subjected to transiently increased cardiomyocyte O-GlcNAc and TAC vs. TAC alone. Additionally, we identified potential transcriptomic markers and driving mechanisms of the exacerbated cardiac pathology. Our findings support a novel concept of cardiomyocyte “O-GlcNAc memory” and suggest key mechanisms of exacerbated cardiac pathology at the transcriptomic level.

## METHODS

### Data Availability

Further materials and reagents used in experiments are found in the Supplemental Materials: Major Resources Table. The data to support findings of this study are accessible upon request from the corresponding author. Files associated with the RNA-sequencing for this study are found on NCBI GEO repository (GSE298401).

### Animal models and experimental design

All studies utilizing mice were done in line with the Institutional Animal Care and Use Committee (IACUC) of the University of Alabama at Birmingham approved protocols (Animal project number: 20530). All procedures involving mice were also compliant with the National Institutes of Health Guide for the Care and Use of Laboratory Animals eighth edition and utilized the ARRIVE reporting system (Supplemental Materials: Major Resources Table & ARRIVE Checklist).

We used the previously generated, validated and characterized dnOGAh mouse.^27,38,49^ In short, the dnOGAh mouse is a model for inducible, cardiomyocyte-specific enhancement of protein O-GlcNAcylation via the cross of previously described mouse α-myosin heavy chain (MHC) promoter driven codon optimized reverse tetracycline trans-activator (αMHC-rtTA, i.e., tON) mice^50^ with previously described tetracycline-responsive element, enhanced green fluorescent protein (EGFP), rat OGA splice variant (TRE-EGFP-dnOGA, i.e., dnOGA) mice.^49^ All study conditions, interventions, groups, and timepoints are outlined in the experimental schema (Figure 1); in more detail, both male and female, 8- to 10-week-old αMHC-rtTA (tON) single transgenic mice were used as controls (Con), and tON + dnOGA double transgenic mice were used as experimental (dnOGAh)—all mice were on the FVB/NJ strain. 2-wk long transgene induction was done via a single intraperitoneal injection of 100 μg doxycycline (DOX) (MilliporeSigma, Burlington, MA, USA) in 0.9 % NaCl solution and switched to 1 g / kg DOX-supplemented NIH31 diet (Envigo, Madison, WI, USA) for 2 weeks. All mice (regardless of Con vs dnOGAh group) were DOX injected and fed DOX chow to control for any indirect effects of DOX treatment.^51^ After 2 weeks of transgene induction, all mice had their cages changed and placed on standard chow for the remainder of the study; 2 weeks after returning to standard chow diet, transverse aortic constriction (TAC) or Sham/Control surgery (described in the next section) was performed on each mouse. Additionally, for all studies, e.g., echocardiography, heart histology, gravimetric measures, molecular studies and left-ventricle (LV) mRNA sequencing, and the mice were climate- and light-control housed at 22°C on a 12 / 12 hours light / dark cycle with *ad libitum* access to food and water throughout the entire study. At the end of the study after 9 weeks of TAC/Sham, mice were anesthetized via 2% isoflurane induction chamber inhalation, then euthanized via manual cervical dislocation followed by organ/tissue excision. Study was conducted in a cohort-matching manner; specifically, cohort-matched mice underwent the dnOGAh ON/OFF-TAC experimental paradigm and were used by cohort for specific measurements/experiments ensuring cohort-matched experimental and control mice for each result; subsequently, mortality/survival data for all mice used in the current study was compiled in the general survival curve analysis (Table S1). To standardize circadian effects,^52-55^ all study terminations and tissue collection was performed between 6-8AM, and all samples were flash frozen with liquid nitrogen and stored at -80°C until subsequent follow up studies.

**Figure 1.**
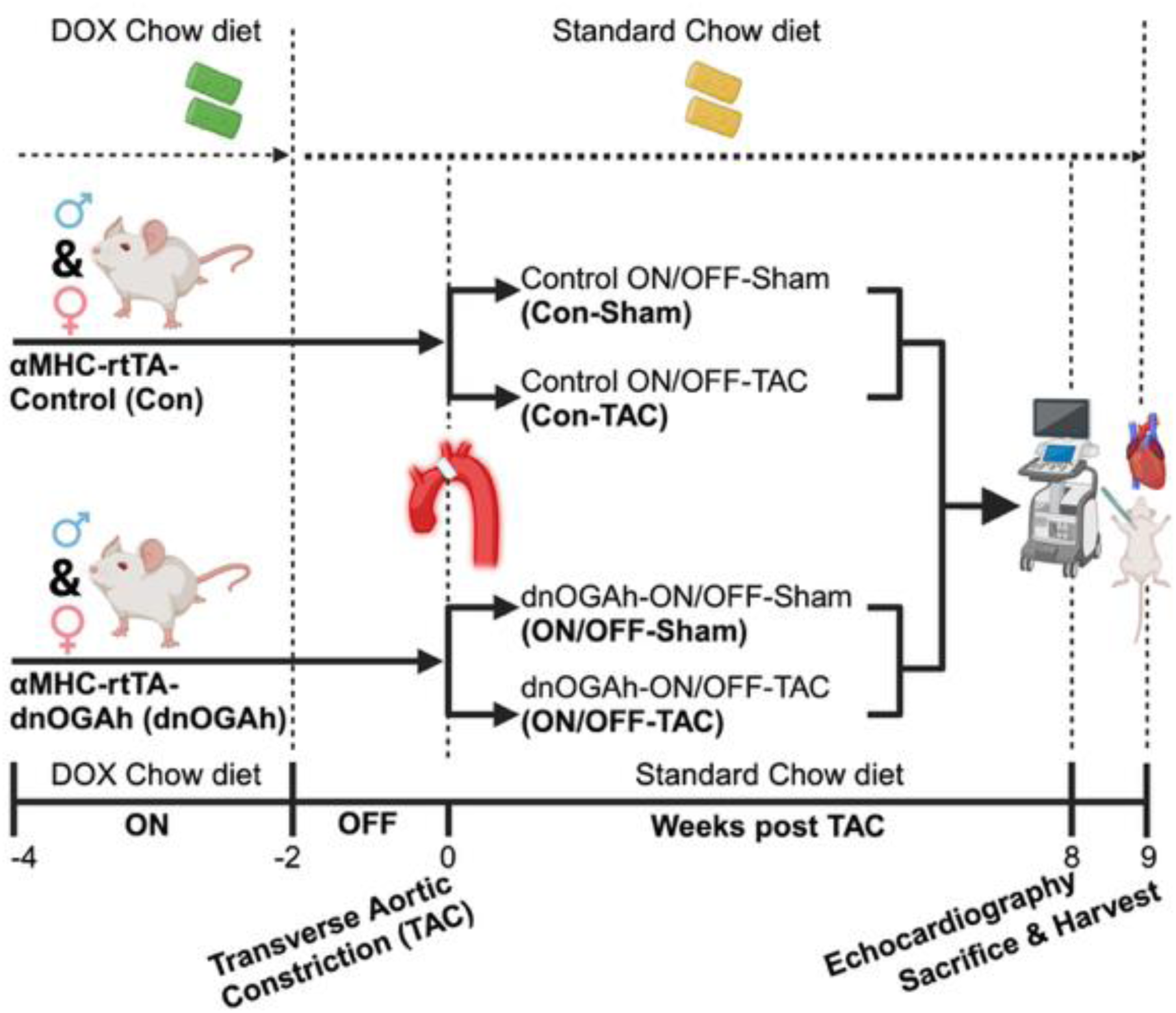
Experimental Schema for Transient Cardiac O-GlcNAc Induction with Subsequent Pressure-Overload (a.k.a., The dnOGAh ON/OFF-TAC Paradigm). Schematic showing the experimental timeline and procedures for the dnOGAh ON/OFF-TAC paradigm used in this study—from induction of dnOGA transgene via DOX, switching the transgene “off” via removal of DOX, and TAC procedure, to subsequent echocardiography and tissue harvesting for downstream analyses.

### Transverse Aortic Construction Surgery

To induce left ventricle pressure overload, we conducted transverse aortic construction (TAC) as previously described;^56-58^ specifically in our study, as shown in the experimental schema (Figure 1), 12- to 14-week old mice (after they underwent 2 weeks on DOX chow and subsequent 2 weeks on standard chow) underwent either TAC or control (Sham) surgery. Mice were anesthetized via 2% isoflurane induction chamber inhalation then placed in a supine position on a heating pad set at 37°C to maintain body temperature; anesthesia was maintained via 1.5% isoflurane and monitored via toe pinching. Fur from neck and chest were removed via topical depilatory cream, and the area was cleaned with ethanol and betadine. At the suprasternal notch, a ∼0.5-1 cm transverse skin incision was made; then followed by a ∼2-3 mm partial superior median sternotomy and the thyroid glands were retracted, to visualize the transverse aortic arch. TAC was induced by placement of a titanium clip (Teleflex Medical, Morrisville, NC, USA)—calibrated to a 30-gauge needle external diameter—on the aorta between the brachiocephalic and left common carotid arteries. After clip placement, chest/sternum and overlying skin were sutured close. Immediately after surgery, mice received intraperitoneal injection of buprenorphine (1 mg/kg) for post-operative pain relief. Sham procedure followed the same steps, but no clip was placed on the aortic arch. Echocardiography, as described below, of the transverse aortic arch at the clip location, and measurement of peak pressure gradient (mmHg) across the constriction location, confirmed TAC induced left ventricle pressure overload following the surgery (Table S2).

### Transthoracic Echocardiography

At the prespecified timepoints shown in the experimental schema (Figure 1) after 8 weeks of TAC/Sham, transthoracic echocardiography was performed on all mice in the study, as previously described.^25,38^ In brief, the mice were anesthetized via 2% isoflurane induction chamber inhalation then placed in a supine position on a heated echocardiography platform (Vevo integrated rail system, VisualSonics) set at 37°C to maintain body temperature; sedation was maintained via nose cone inhalation of 1.0-1.5% isoflurane which allowed for heart rate maintenance of ∼400 beats / min. Fur from the chest was removed via topical depilatory cream, and then the mice were then placed in a left-lateral-decubitus position. Image acquisition was obtained on a Vevo 770 high-resolution in vivo micro-imaging system (VisualSonics) using an RMV-707B 30MHz ultrasound probe (VisualSonics); subsequently, images were analyzed on the Vevo 770 software using the Cardiac Measurements Package (VisualSonics). This software and package allowed for measurement and calculation in M-mode images of various cardiac parameters such as Ejection Fraction (EF), Stroke Volume (SV), Fractional Shortening (FS) and Cardiac Output (CO); additionally, measurements such as LV internal dimension (LVID) and posterior wall (LVPW) thickness, and Interventricular septal thickness (IVS) at both diastole (;d) and systole (;s).

### Western Blotting

Detection and analysis of specific proteins and pan-protein O-GlcNAcylation was done via western blotting as previously described and optimized.^27,38,59^ In brief, total LV tissue was homogenized and protein extracted using Radioimmunoprecipitation assay (RIPA) buffer consisting of 150 mM NaCl, 1 mM EDTA, 1 mM EGTA, 50 mM Tris-pH 7.5, 1% NP-40, 1% sodium deoxycholate, 1 mM Na_3_VO_4_, 10 μM Thiamet-G (SML0244, MilliporeSigma), Halt protease/phosphatase inhibitor cocktail (Thermo Fisher Scientific, Waltham, MA, USA), and 10 μM phenylmethylsulfonyl fluoride (PMSF) (36978, ThermoScientific). 20-30 μg. of protein lysate was loaded in each lane and separated on 8 or 12% sodium dodecyl sulfate (SDS) polyacrylamide gels, then transferred to a 0.45 μm immobilon-FL PVDF membrane (IPFL00010, MilliporeSigma); note that the transfer of protein O-GlcNAcylation blots used transfer buffer containing 0.375% SDS, as described previously.^59^ Membranes were immunoblotted using primary antibodies for specific proteins—OGA (14711-1-AP, ProteinTech), OGT (O6264, MilliporeSigma), and β-actin (GTX629630, GeneTex), β-Tubulin (ab131205, Abcam)—in 1X casein blocking buffer (B6429, MilliporeSigma). Fluorescence conjugated secondary antibodies—either goat anti-rabbit IgG (A21109, ThermoScientific) or donkey anti-mouse IgG (936-32212, Li-Cor Biosciences, Lincoln, NE, USA)— were used for protein detection, and membranes were scanned on the Odyssey CLx infrared imaging system (9120, Li-Cor Biosciences). For protein O-GlcNAcylation, membranes were immunoblotted using a primary antibody—CTD110.6 (UAB Epitope Recognition and Immunoreagent Core)—in 3% Bovine Serum Albumin (BSA) blocking buffer (sc-2323, Santa Cruz Biotechnology). Protein detection used secondary goat anti-mouse IgM-HRP (401225, Calbiochem) and SuperSignal West Pico PLUS Chemiluminescent Substrate (34580, ThermoScientific), and was captured on an Amersham Imager 600 (GE Health Care, Chicago, IL, USA). All western blot images were analyzed on Image Studio Lite (Li-Cor Biosciences).

### Histological Staining and Analysis

Histological studies were performed as previously described.^38^ In brief, medial, transverse LV tissue sections were fixed in 10% formaldehyde (F8775, Millipore-Sigma) and embedded in paraffin. Subsequent histology sectioning, slide mounting, and staining (for both Hematoxylin & Eosin (H&E) and Trichome stains) were performed at the UAB Comparative Pathology Laboratory. Staining was confirmed and images captured via light microscopy (Nikon Eclipse E200 with NIS element software); both quantification of trichrome signal, H&E signal, and measurement of cross-sectional cardiomyocyte cell area was performed in ImageJ Software (NIH).

### RNA-Isolation for Sequencing and mRNA Sequencing Analysis

Bulk RNA sequencing was performed on whole LV tissue from both male and female mice in each of the four experimental conditions, as shown in the experimental schema (Figure 1; n = 5 per sex, per group). RNA was isolated from LV tissue using the RNeasy® Fibrous Tissue Mini Kit (Qiagen Inc.), following the manufacture’s protocol. To ensure quality, isolated RNA was analyzed to have an RNA integrity number of at least 7. Subsequent library preparation, quality control, and sequencing steps were formed by the UAB Heflin Center for Genomic Studies, Genomics Core Laboratory; next-generation RNA sequencing was performed on an NovaSeq 6000 system (Illumina), generating paired-end 100-bp reads. Full details on the upstream processing and downstream analyses of the RNA sequencing data for this study can be found in the associated GitHub repository https://github.com/samuelfchang/dnOGA_ONOFF_TAC_RNA_Seq_Proj. In brief, upstream processing was performed on the UAB IT Research Computing, High Performance Computing fabric (a.k.a., “Cheaha”), referencing the nf-core pipeline for RNA sequencing upstream processing (nf-core/rnaseq v3.18.0);^60^ specifically, aligned to mouse genome GRCm39 (GENCODE release M32). Downstream analyses and visualization were performed using R (v4.3.1), in R-Studio (v2024.12.0+467), with differential gene expression analysis performed using and following documentation for *DESeq2* (v1.40.2),^61^ and as previously described.^38^ Specifically, using *DESeq2*, differential gene expression quantification and significance testing was determined with the Wald test which output: quantile-normalized read counts (via negative binomial generalized linear model fitting), log_2_ fold change for each contrast (changes in gene expression in pairwise comparisons between each of the experimental groups), p-values, and Benjamini-Hochberg adjusted p-values (False Discovery Rate, or FDR/q-values). Further analyses and data visualization came from importing these results into: VennPlex^62^ to compare and visualize differentially expressed genes (DEG) of the various contrasts (DEG filtering cutoff: q <0.05 & Fold Change (FC) ≥ |1.5|), STRING database (STRING-db)^63^ for gene level associated protein-protein interaction networks and pathway analysis (STRING-db parameters: Network type-Full STRING network, Required score-medium confidence at 0.400, FDR stringency < 0.05) for DEGs (filtering cutoff: q <0.05 & FC ≥ |1.5|), and Pathview^64^ using KEGG^65-67^ data for biological pathway analysis and visualization for DEGs (q < 0.05).

Files for the RNA-sequencing in this study are found on the NCBI GEO repository (GSE298401); and all details on the upstream processing and downstream analyses (e.g., packages, parameters, and code) for the RNA sequencing data in this study can be found in the associated GitHub repository: https://github.com/samuelfchang/dnOGA_ONOFF_TAC_RNA_Seq_Proj.

### Statistical Analysis

Excluding the RNA Sequencing data, statistical analyses were performed using GraphPad Prism 10.4.1 on MacOS X (GraphPad Software). Tests performed included: two-way ANOVA followed by Tukey’s multiple comparisons test. Unpaired non-parametric Mann-Whitney *t* test (2-tailed). Results are presented as Mean ± S.E.M., with cutoff for statistical significance set at p < 0.05. For survival analysis, survival curves were analyzed via Log-Rank (Mantel-Cox) Test. Results are presented as X^2^ and degrees of freedom, with cutoff for statistical significance set at p < 0.05. For quantification of exacerbated pathology/change in measurements, Bliss independence calculation for synergy was used.^68^ Results are presented as combination index (*CI*) with cutoff for synergy set at *CI* < 1. For RNA sequencing data, as mentioned earlier, using *DESeq2*, differential gene expression quantification and significance testing was determined with the Wald test. Specific gene normalized read count (normalized counts) results were graphed in GraphPad Prism and presented as Mean ± S.E.M.; and statistical significance was pulled from the *DESeq2* Wald Test results for p < 0.05, q < 0.1, and q < 0.05.

### Graphics Generation

Experimental Schema (Figure 1) and elements of the graphical abstract were generated using BioRender Software (BioRender, Toronto, Ontario, Canada).^69,70^

## RESULTS

### Transient Cardiomyocyte-specific O-GlcNAc Induction (dnOGA ON/OFF) with Subsequent Pressure Overload and Levels of Cardiac Total Protein O-GlcNAc

We previously reported the dnOGA transgene is specifically expressed in hearts of tON-dnOGA, double transgenic, mice when DOX is present,^27^ and that cardiomyocyte dnOGA expression leads to a significant increase in cardiac protein O-GlcNAcylation at 2 weeks of dnOGA induction.^27,38^ We confirm these finding here demonstrating DOX induces the dnOGA transgene with a corresponding significant increase in protein O-GlcNAcylation at 2 weeks of induction (Figure S1C-S1F). However, we observe no significant difference in cardiac protein O-GlcNAcylation after 2 weeks back on standard chow, or across the four experimental groups at the end of the dnOGAh ON/OFF-TAC paradigm (Figure 2A, 2B, S1A, S1B, S2A-S2C). OGA levels (Figure 2C and S2E) do not change after 2 weeks back on standard chow or between any of the experimental end groups. OGT levels only modestly increase in the Con-TAC vs. Con-Sham comparison (Figure 2D). Additionally, we observed no differences in survival curves between female and male mice, as well as between the four experimental groups (Table S1). Taken together, these results show that in this model of transiently enhanced cardiomyocyte protein O-GlcNAc and subsequent pressure-overload achieves a temporary increase in cardiac protein O-GlcNAcylation with a subsequent return to control levels of cardiac protein O-GlcNAcylation prior to TAC or Sham surgery.

**Figure 2.**
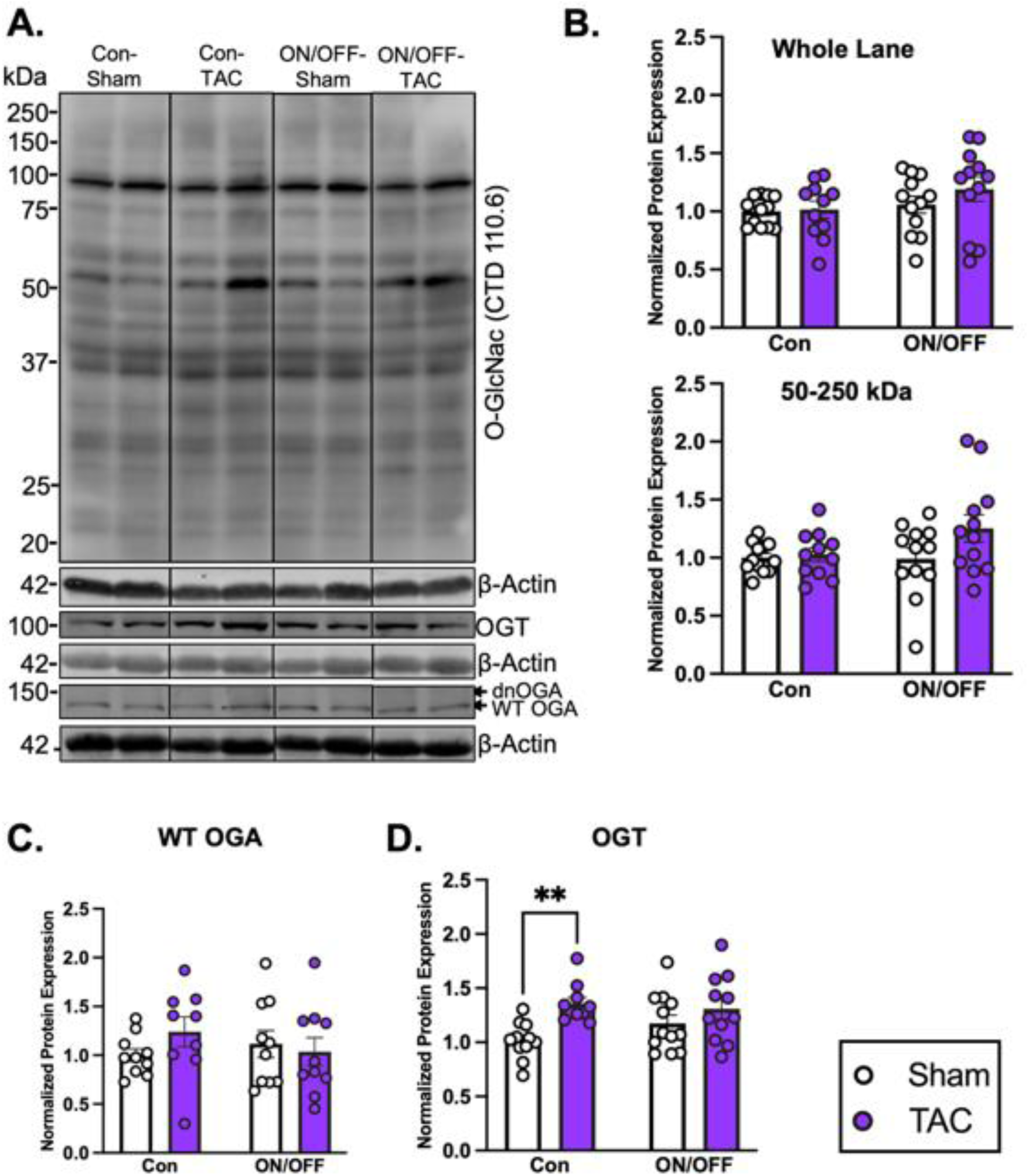
Transient cardiomyocyte-specific O-GlcNAc induction (dnOGA ON/OFF) with/without pressure overload shows no change in cardiac total protein O-GlcNAc levels compared to respective controls: a validation of the dnOGAh ON/OFF model. **A**) Western blot assay of left ventricle (LV) whole cell protein extract for protein O-GlcNAcylation (via CTD110.6), protein levels of endogenous/WT OGA (lower band), dnOGA (upper band due to eGFP fusion), and OGT for Con-Sham, Con-TAC, ON/OFF-Sham and ON/OFF-TAC conditions. **B-D**) Densitometry analysis of all immunoblots. **B**) Top panel: quantification of whole-lane cardiac protein O-GlcNAcylation (20–250 kDa protein size). Bottom panel: quantification of cardiac protein O-GlcNAcylation at 50–250 kDa protein size. **C**) quantification of endogenous/WT OGA protein. **D**) quantification of OGT. All quantitative data are mean ± S.E.M (*n* ≥ 9); Statistical significance was based on two-way ANOVA followed by Tukey’s multiple comparisons test (** *p* < 0.01).

### Transient Cardiac O-GlcNAc Induction with Subsequent Pressure-Overload induces Cardiac Hypertrophy, Fibrosis and Dysfunction

As expected, total-heart weight, LV-weight, and LV cross-sectional cardiomyocyte cell area increased significantly in the Con-TAC and ON/OFF-TAC vs. respective Sham conditions (Figure 3A-3C and 3E), with no change of RV weight in any group (Figure S3A). Importantly, total-heart weight, LV-weight and LV cardiomyocyte cell area all show further significant increase in ON/OFF-TAC vs. Con-TAC groups—with no significant differences in any index of hypertrophy between ON/OFF-Sham vs. Con-Sham groups.

**Figure 3.**
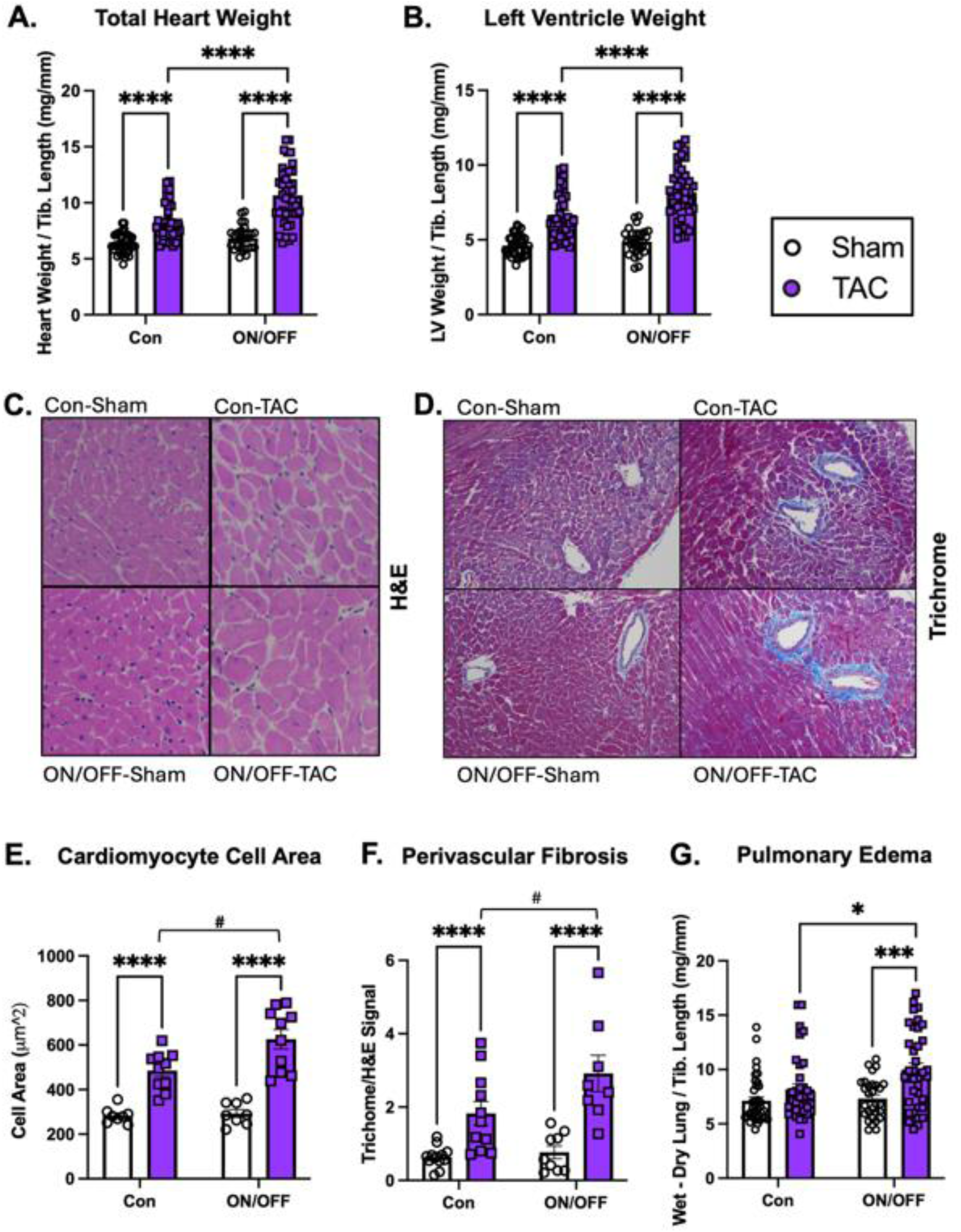
Transient cardiac O-GlcNAc induction with subsequent pressure-overload induces further cardiac hypertrophy, perivascular fibrosis and dysfunction compared to pressure-overload alone. **A)** Normalized ratio of total heart weight to tibia length (TL), and **B**) left ventricle (LV) plus septum to TL, for Con-TAC and ON/OFF-TAC mice compared to age-matched Sham controls (Con-Sham and ON/OFF-Sham) (*n* ≥ 27). **C-D**) Representative images from histological examination of LV cardiac tissue using hematoxylin and eosin (H&E) staining (**C**), and perivascular LV cardiac tissue H&E and trichrome staining (**D**). **E)** Average cross-sectional LV cardiomyocyte cell area for Con-TAC and ON/OFF-TAC mice compared to age-matched Sham controls (Con-Sham and ON/OFF-Sham) (*n* ≥ 7). **F)** Normalized ratio of trichrome to H&E signal of perivascular LV tissue for Con-TAC and ON/OFF-TAC mice compared to age-matched Sham controls (Con-Sham and ON/OFF-Sham) (*n* ≥ 8). **G)** Normalized ratio of wet-dry lung weight to TL for Con-TAC and ON/OFF-TAC mice compared to age-matched Sham controls (Con-Sham and ON/OFF-Sham) (*n* ≥ 29). Results for previous measures split by sex are found in Figure S3. Statistical significance was based on: two-way ANOVA followed by Tukey’s multiple comparisons test (**A**–**G**) (* *p* < 0.05, *** *p* < 0.01, **** *p* < 0.001), unpaired non-parametric Mann-Whitney (two tailed, **E-F**) (# *p* < 0.05). All calculated and analysis data are mean ± S.E.M.

Given the known association between pathologic cardiac hypertrophy and fibrosis,^71,72^ we evaluated LV tissue fibrosis across the four experimental groups. Using histologic measurement of trichrome staining in LV tissue, perivascular fibrosis significantly increased in both Con-TAC and ON/OFF-TAC groups compared to their respective Sham controls (Figure 3D and 3F). Moreover, perivascular cardiac fibrosis further increases in ON/OFF-TAC vs. Con-TAC groups. Meanwhile, interstitial fibrosis showed no significant change between the four groups (Figure S4F, S4G).

With results showing further increases in both LV hypertrophy and peri-vascular fibrosis, we assessed cardiac function across the four experimental groups. Pulmonary edema as a measure of cardiac dysfunction showed no significant change in the Con-TAC vs. Con-Sham comparison, but importantly, significantly increased in the ON/OFF-TAC vs. ON/OFF-Sham and ON/OFF-TAC vs. Con-TAC comparisons (Figure 3G); this showed pulmonary edema further increased significantly in ON/OFF-TAC vs. ON/OFF-Sham groups compared to Con-TAC vs. Con-Sham groups. Measures of cardiac function via echocardiography show expected decreases in ejection fraction (EF) in both Con-TAC and ON/OFF-TAC groups compared to their respective Sham controls (Figure S5A), but no further decrease in the ON/OFF-TAC vs. Con-TAC groups

Taken all together, across the various cardiac hypertrophy, fibrosis, and function measures, transiently increased cardiac protein O-GlcNAcylation with subsequent pressure-overload further significantly increased cardiac pathologic remodeling compared to pressure-overload alone. Specifically, results show further increases in LV cardiomyocyte hypertrophy, perivascular fibrosis, and cardiac dysfunction—not due to further impaired systolic function.

### Transcriptomic Analyses of Transiently Increased Cardiac Protein O-GlcNAc with Subsequent Pressure-Overload both Confirm Markers and Identify Novel-Pathways of Exacerbated Pathologic Cardiac Remodeling

In light of our earlier work demonstrating the effect of sustained increased cardiomyocyte protein O-GlcNAcylation on the LV transcriptome, as well as the known role of O-GlcNAc to regulate transcriptional and epigenetic regulation,^31,38^ we performed mRNA sequencing to identify and examine the impact of transiently increased cardiomyocyte protein O-GlcNAc with subsequent TAC on the global LV transcriptome. After relevant quality control, filtering, and normalization (Figure S6A-S6I), unsupervised principal component analysis (PCA) showed distinct transcriptomic profiles, separated by principal component 1, of each of the four experimental groups (Figure S6H); filtering for differentially expressed genes (DEGs) shown via Volcano plot (Figure S7A), and frequency tables of various significance cutoffs (Figure S7B). Full tables of DEGs at various significance cutoffs are found in Table S3, with key in Table S4. Candidate search for genes related to our phenotypic results in previous figures show that transcriptomic markers for cardiac hypertrophy, fibrosis and dysfunction all follow the pattern of significant further changes in the ON/OFF-TAC vs. Con-TAC comparison; for instance, related to cardiac structure and hypertrophic remodeling, *My6* (α**-**myosin heavy chain) expression significantly decreases in the ON/OFF-TAC vs. Con-TAC comparison (Figure S8A). With regards to fibrosis, *Col1a1* (Collagen type 1 α-1) expression across the four groups recapitulates the changes we show in peri-vascular fibrosis (Figure 3D and 3F) and shows further significant increase in expression shown in the ON/OFF-TAC vs. Con-TAC groups (Figure S8B). Also, *Nppa* (Atrial natriuretic peptide, ANP) expression, as a marker of cardiac dysfunction, shows further significant increase in expression shown in the ON/OFF-TAC vs. Con-TAC comparison (Figure S8C). Additional gene markers of pathologic cardiac remodeling follow similar patterns of significant change (Figure S8D-S8I). Taken as a whole, the expression changes in these specific gene markers transcriptomically support our previous phenotypic results of furthered pathologic cardiac remodeling.

Beyond transcriptomic confirmation of furthered pathology, subsequent analysis of DEGs (filtered at FDR/q-value < 0.05 and |FC| ≥ 1.5) via Venn analysis uncovers potential drivers and effectors of the furthered pathology we observe in our phenotypic results. Venn analysis resolves DEGs associated with various overlapping comparisons between the four experimental groups (Figure 4A), and we specifically focus on DEGs associated with the key comparison of further changed expression in the ON/OFF-TAC vs. Con-TAC groups; a table of what each Venn region represents in terms of significant changes in the various comparisons are shown in Table S5, with all DEGs from the 4-way Venn diagram found in Table S6. To identify transcriptomic pathways and regulatory mechanisms within the DEGs of the Venn analysis, we perform pathway enrichment following network interaction analyses using STRING-db on the up- and down-regulated DEGs from the key ON/OFF-TAC vs. Con-TAC comparison.

**Figure 4.**
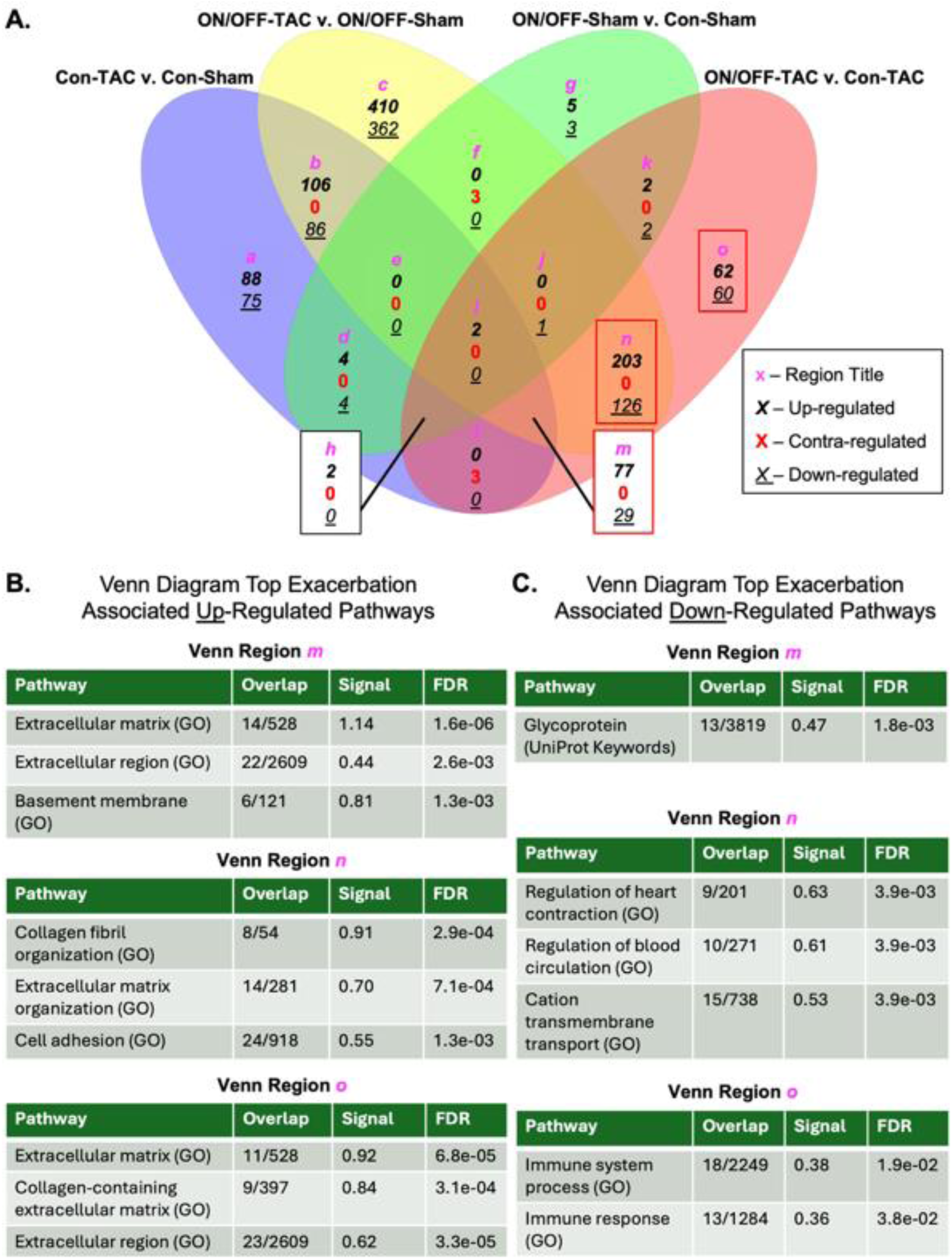
Venn diagram of DEGs from relevant pairwise comparisons and pathway analyses of DEGs from key Venn regions reveal transcriptomic markers of pathology exacerbation. **A)** Venn diagram of DEGs from four comparisons between experimental groups; table of all DEGs found in the various regions of the Venn diagram are found in Supplemental Table 6 (Table S6 in the Data Supplement). **B-C)** Pathway enrichment analysis of DEGs utilizing STRING-db with Gene-Ontology (GO) and UniProt Keyword databases; top significantly enriched pathways for upregulated DEGs in regions (**B**), and top significantly enriched pathways for downregulated DEGs in regions (**C**). Table of all enriched pathways and regulated genes within each are found in Supplemental Table 7 (Table S7 in Data Supplement).

In Figure 4B-4C, we show significantly enriched pathways for three important Venn diagram regions associated with furthered significant changes in the ON/OFF-TAC vs. Con-TAC comparison. The top upregulated pathways for all three regions associate with ECM and fibrosis pathways (Figure 4B): examples include, Extracellular matrix (GO0031012), Collagen-containing extracellular matrix (GO0062023), Basement membrane (GO0005604), Collagen fibril organization (GO 0030199), Cell adhesion (GO 0007155), and Extracellular region (GOCC0005576). Top downregulated pathways across the three regions include (Figure 4C): Glycoprotein (UniProt KW-03525, protein containing one or more covalently linked carbohydrates of various kinds), regulation of heart contraction (GO), regulation of blood circulation (GO), cation transmembrane transport (GO), Immune system process (GO) and Immune response (GO). All significantly enriched pathways from the DEGs of the Venn diagram can be found in Supplemental Table 7 (Table S7). From the pathway analysis of DEGs from key regions in the 4-way Venn analysis, transcriptomic signatures identify biological/molecular pathways which could potentially drive the furthered cardiac pathology seen in O-GlcNAc memory with subsequent pressure-overload.

### DEGs from Pathway Analysis and Candidate DEGs Identify Specific Transcriptomic Drivers of Pathology Exacerbation From Transiently Increased Cardiac Protein O-GlcNAc with Pressure-Overload

Examining the individual DEGs which make up the significantly enriched up- and down-regulated pathways in Figure 4, we find key DEGs that encode proteins associated specifically within the context of furthered cardiac pathology. In the enriched upregulated pathways associated with ECM/fibrosis/collagen, we find upstream “driver” regulatory mechanisms for ECM associated pathologic cardiac fibrotic remodeling (Figure 5A, 5B and S9A). Examples include: *Ccn2* (Connective tissue growth factor, CTGF, Figure 5A), and *Postn* (Periostin, Figure 5B); in both DEGs, the Con-TAC vs. Con-Sham and ON/OFF-TAC vs. ON/OFF-Sham comparisons show significantly increased gene expression, and importantly, the key ON/OFF-TAC vs. Con-TAC comparison shows significantly further increased expression.

**Figure 5.**
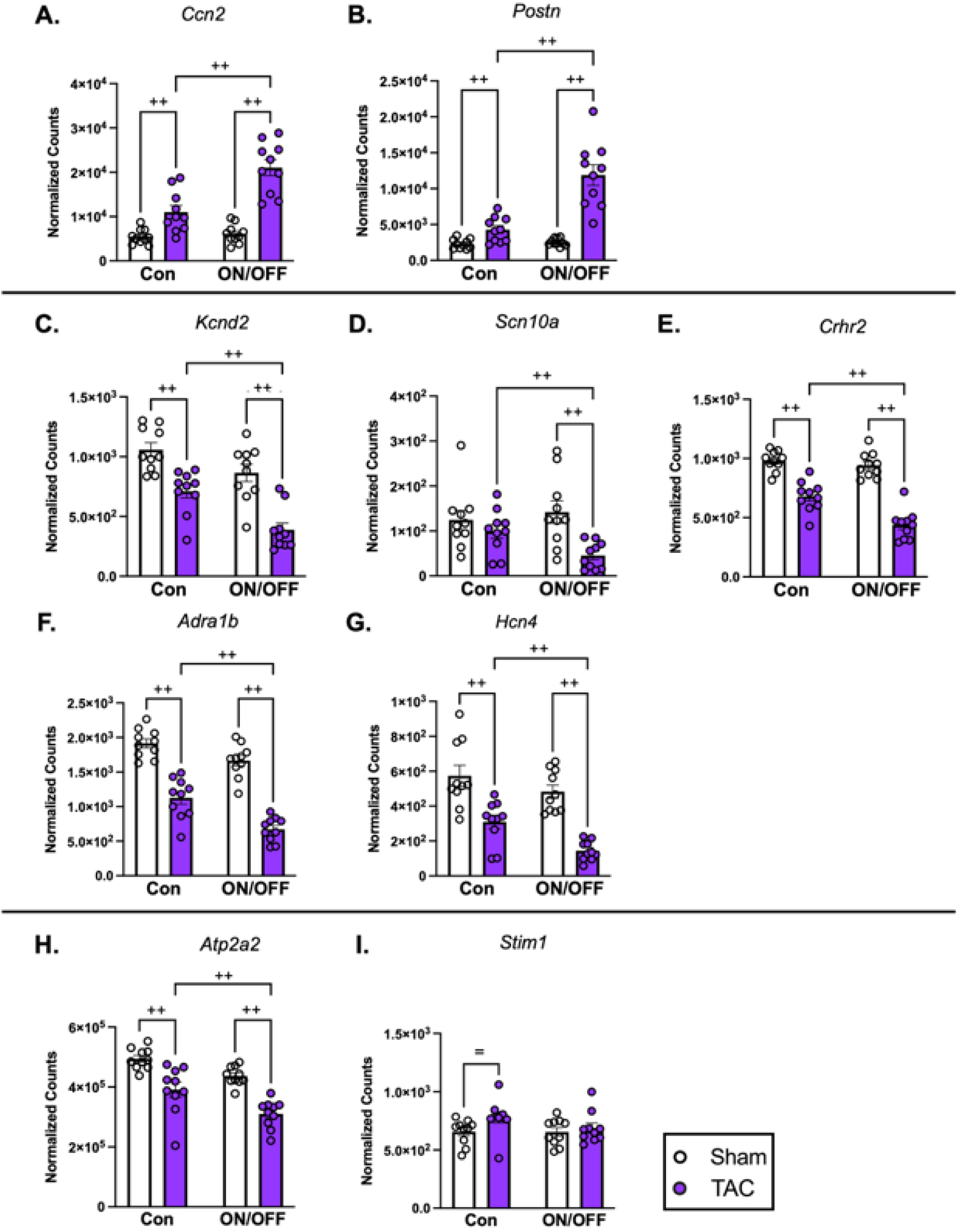
DEGs from pathway analyses of key Venn regions and candidate DEGs identify potential transcriptomic drivers of pathology exacerbation. **A-I)** Specific DEGs from mRNA sequencing of LV tissue (*n* = 5 per sex, per group) in normalized counts that were significantly enriched in pathway analysis of Figure 7. **A-B**) Specific DEGs from pathways associated with extracellular matrix/fibrosis/collagen: Connective tissue growth factor (CTGF) (*Ccn2*), and Periostin (*Postn*). **C-E**) Specific DEGs from pathways associated with heart contraction/function and ion transport: potassium channels (*Kcnd2*), subunit of a voltage-gated sodium channel (*Scn10)*, and corticotropin-releasing hormone receptor 2 (*Crhr2*). **F-G**) Specific DEGs from pathways associated with glycosylated proteins: β-1A adrenergic receptor for catecholamine binding (*Adra1b*), and hyperpolarization-activated cyclic nucleotide-gated potassium channel 4 (*Hcn4*). **D-I**) Specific DEGs associated with calcium signaling: sarcoplasmic/endoplasmic reticulum Ca^2+^-ATPase 2a (SERCA2a) (*Atp2a2*), and stromal interaction molecule 1 (*Stim1*). Statistical significance was based on Wald test (**A-I**) (*= p* < 0.05, + *q* < 0.1, and ++ *q* < 0.05). All calculated and analysis data are mean ± S.E.M.

Investigating DEGs from the significantly enriched downregulated pathways associated with heart contraction/function and ion transport, the majority of key DEGs encode proteins associated with some form of cardiac ion transport (Figure 5C-5E, and S9B): for example, *Kcnd2* encodes a potassium channel (Figure 5C), and *Scn10a* encodes the subunit of a voltage-gated sodium channel (Figure 5D). And other specific DEGs include genes which encode receptor proteins for hormones which can influence cardiac function/rhythm, such as *Crhr2* (corticotropin-releasing hormone receptor 2, Figure 5E). Similarly, although significantly enriched in a different pathway associated with glycosylated proteins, key DEGs from this downregulated pathway also primarily encode proteins associated with either ion transport or hormones impacting cardiac function/rhythm (Figure 5F, 5G and S9C): *Adra1b* (β-1A adrenergic receptor for catecholamine binding, Figure 5F), and *Hcn4* (hyperpolarization-activated cyclic nucleotide-gated potassium channel 4, Figure 5G).

Observing the trend of key ion transport DEGs from the downregulated pathway analyses, we perform a candidate search for DEGs associated with calcium handling/signaling based on our previous publication (Figure 5H, 5I and S9D).^38^ Some examples include: *Atp2a2* encoding Sarcoplasmic/endoplasmic reticulum Ca^2+^-ATPase 2a (SERCA2a, Figure 5H). And *Stim1* encoding known O-GlcNAcylation target stromal interaction molecule 1 (STIM1)^73^ is only upregulated in the Con-TAC vs. Con-Sham comparison (Figure 5I). Returning to the pathway analysis, investigating the DEGs from the significantly enriched downregulated pathways also shows some genes associated with immune system processes. Taken as a group, these DEGs appear to be generally associated with B-cell immunity (Figure S9E); for example, *Bcl11a* (B-cell lymphoma/leukemia 11a), and *Shld1* (Shieldin complex subunit 1) which plays a role in immunoglobulin class-switch recombination and DNA-repair in B-cells.^74^ Taken all together, filtering and analysis of key individual DEGs from pathway analyses identifies cardiac relevant genes, encoding proteins associated with key biological processes in the heart, which could indicate transcriptionally upstream drivers of the furthered cardiac pathology we show in previous figures.

### Transiently Increased Cardiac Protein O-GlcNAc with Subsequent Pressure-Overload Causes Transcriptomic Changes Associated with Altered Redox Mechanisms, Cardiac Metabolism and Mitochondria

Previous studies demonstrate a clear link between increased cardiac protein O-GlcNAc and decreased mitochondrial function;^25,27,75^ specifically, our previous publication on sustained cardiomyocyte protein O-GlcNAcylation showed not only transcriptomic and proteomic changes in redox mechanisms and suppressed energy metabolism/mitochondrial pathways in the heart, but also, reduced oxidative phosphorylation (OXPHOS) mitochondrial function.^38^ In our current study, with regards to changes in redox mechanisms, *Nox4* (NADPH-Oxidase 4, NOX4) expression significantly increases in the ON/OFF-TAC vs. ON/OFF-Sham comparison while non-significantly changed in Con-TAC vs. Con-Sham comparison; importantly, *Nox4* expression significantly increases in the key ON/OFF-TAC vs. Con-TAC comparison (Figure 6A). NOX4 is a cardiac reactive oxygen species (ROS) producer that is heavily implicated in the cardiac response to metabolic syndrome and/or pressure overload stressors.^76-83^ Another isoform of NOX, *Cybb* encoding NOX2, only significantly increases in the ON/OFF & TAC comparison (Figure S10A). Conversely, mitochondrial antioxidant process gene *Sod2,* Superoxide dismutase 2 mitochondrial, expression significantly decreases in the ON/OFF-TAC vs. ON/OFF-sham comparison and key ON/OFF-TAC vs. Con-TAC comparison (Figure 6B); furthermore, expression of key regulator of redox pathways and known O-GlcNAcylation target,^84^ *Nfe2l2* encoding Nuclear factor erythroid 2-related factor 2 (NRF2), increases only in the Con-TAC vs. Con-Sham comparison—at raw p-value < 0.05 (Figure 6C).

**Figure 6.**
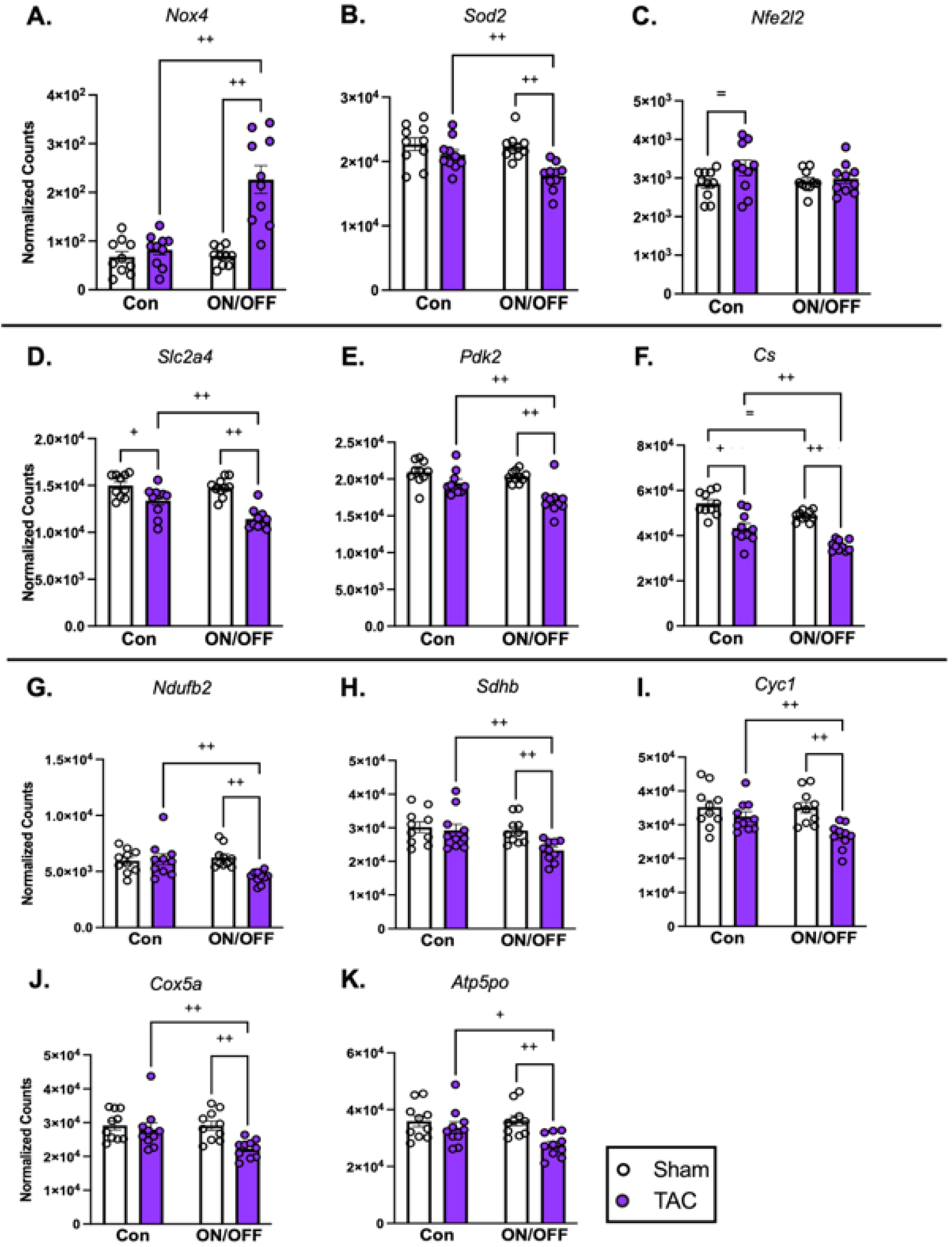
Transient Cardiac O-GlcNAc induction with subsequent pressure-overload changes transcriptomic profiles associated with redox mechanism, and cardiac metabolism. **A-K)** Specific candidate DEGs from mRNA sequencing of LV tissue (*n* = 5 per sex, per group) in normalized counts. **A-C**) Specific DEGs associated with redox mechanisms: NADPH-Oxidase 4 (*Nox4*), superoxide dismutase 2 mitochondrial (*Sod2*), Nuclear factor erythroid 2-related factor 2 (*Nfe2l2*). **D-F**) Specific DEGs associated with cardiac metabolism: glucose transporter GLUT4 (*Slc2a4*), Pyruvate dehydrogenase kinase 2 (*Pdk2*), Citrate synthase (*Cs*). Other DEGs associated with the citric acid cycle are found in Figure S11. **G-K**) Specific DEGs associated with the electron transport chain: NADH:ubiquinone oxidoreductase subunit B2 of Complex I (*Ndufb2*), succinate dehydrogenase complex subunit B of Complex II (*Sdhb*), Cytochrome c1 of Complex III (*Cyc1*), Cytochrome c oxidase subunit 5A of Complex IV (*Cox5a*), ATP synthase subunit O of Complex V (*Atp5po*). Other DEGs associated with the electron transport chain are found in Figure S12. Statistical significance was based on Wald test (**A-K**) (*= p* < 0.05, + *q* < 0.1, and ++ *q* < 0.05). All calculated and analysis data are mean ± S.E.M.

In terms of genes associated with metabolic processes, we first explored key DEGs encoding proteins involved in glucose metabolism. *Slc2a4* expression, encoding the predominant cardiac glucose transporter GLUT4, significantly decreased in both Con-TAC vs. Con-Sham and ON/OFF-TAC vs. ON/OFF-Sham comparisons, and importantly significantly further decreased in the ON/OFF-TAC vs. Con-TAC comparison (Figure 6D). Linking glycolysis and mitochondrial glucose oxidation, *Pdk2* (Pyruvate dehydrogenase kinase 2) expression significantly decreases in both the ON/OFF-TAC vs. ON/OFF-Sham and ON/OFF-TAC vs. Con-TAC comparisons (Figure 6E); however, both the *Pdk1* and *Pdk4* isoforms are not significantly changed in any comparison (Figure S10B). Furthermore, investigating the genes encoding enzymes of the tricarboxylic acid (TCA) cycle, *Cs* (citrate synthase) expression is significantly decreased in all four comparisons—although ON/OFF-Sham vs. Con-Sham comparison is only via raw p-value (Figure 6F); and generally, other TCA cycle genes exhibit similar significantly decreased expression across a combination of the four comparisons (Figure S11).

Moving to mitochondrial OXPHOS complex associated genes, select DEGs representative of each of the OXPHOS complexes show similar significant downregulation in both ON/OFF-TAC vs. ON/OFF-Sham and the ON/OFF-TAC vs. Con-TAC comparisons: *Ndufb2* (encoding NADH:ubiquinone oxidoreductase subunit B2 of Complex I, Figure 6G), *Sdhb* (encoding succinate dehydrogenase complex subunit B of Complex II, Figure 6H), *Cyc1* (encoding Cytochrome c1 of Complex III, Figure 6I), *Cox5a* (encoding Cytochrome c oxidase subunit 5A of Complex IV, Figure 6J), and *Atp5po* (encoding ATP synthase subunit O of Complex V, Figure 6K). To provide a wider view, DEGs—for the ON/OFF-TAC vs. Con-TAC comparison (at q < 0.05) imported into KEGG Pathview—return a global OXPHOS visualization of associated gene regulation (Figure S12); like our candidate approach (Figure 6G-6K), the KEGG pathway visualization shows global decreased gene expression across OXPHOS complex associated genes in the ON/OFF-TAC vs. Con-TAC comparison.

With all the changes in metabolism associated gene expression, we investigated regulatory processes of mitochondrial homeostasis and oxidative capacity. First, *Ppargc1a* expression (encoding PPARG coactivator 1 alpha, PGC-1α), a key transcriptional coactivator and regulator of mitochondrial biogenesis,^85^ significantly decreased in both the ON/OFF & TAC and ON/OFF-TAC vs. Con-TAC comparisons (Figure S10C); interestingly, the β isoform (*Ppargc1b/*PGC-1β) expression decreased in the ON/OFF-sham vs. Con-Sham and ON/OFF-TAC vs. Con-TAC comparisons (Figure S10C).

Finally, as the hexosamine biosynthetic pathway (HBP), which synthesizes the PTM residue for O-GlcNAcylation—UDP-GlcNAc, also branches off from glycolysis, we investigated whether O-GlcNAc memory with subsequent pressure-overload affects HBP genes; *Gfpt1* expression, encoding the HBP rate-limiting enzyme glutamine-fructose-6-phosphate transaminase 1 (GFAT1), does not significantly change in any of the comparisons (Figure S10D), and *Uap1* (UDP-N-acetylglucosamine pyrophosphorylase) expression levels only significantly decrease in the ON/OFF-TAC vs. Con-TAC comparison (Figure S10D).

Taking all the candidate DEG analysis together, our results show that O-GlcNAc memory with subsequent pressure-overload further changes gene expression in the key ON/OFF-TAC vs. Con-TAC comparison associated with redox mechanisms, metabolism and mitochondrial homeostasis.

## DISCUSSION

The current study tested the hypothesis that a transient increase in cardiomyocyte protein O-GlcNAcylation leads to exacerbated adverse cardiac remodeling following subsequent pressure overload. The overall rationale for this study is based in the results of the DCCT/EDIC studies, which found a significant increase in MACE rates for diabetic patients on conventional glycemic management during DCCT then switched to tight glycemic control therapy during EDIC vs. patients always on tight glycemic control therapy (i.e., goal HbA1c at near-normal ranges);^9-12^ these results, along with studies such as the UKPDS and VADT for type-2 diabetes, introduce the concept of “metabolic memory” and how it affects long-term outcomes for diabetic patients.^14-17^ Potential genetic/molecular mechanisms underlying metabolic memory include epigenetic modifications—for example, changes in DNA methylation status due to previous hyperglycemia.^18,19^ Recent research on cardiac O-GlcNAcylation in the context of CVD and diabetes lays the scientific foundation for investigation O-GlcNAc as a key factor in the development of diabetic cardiac pathology. For example, Prakoso et. al. showed not only increased cardiac O-GlcNAc in diabetic patients vs. non-diabetic patients, but also a significant positive correlation between increasing cardiac O-GlcNAc and LV dysfunction.^28^ Furthermore, our recent work studying effects of sustained cardiomyocyte O-GlcNAc enhancement recapitulates many hallmarks of pathology seen in diabetic hearts.^38^ Related to potential mechanisms of metabolic memory, O-GlcNAc is also known to play a role in both transcriptional and epigenetic regulation, and can affect cellular memory;^31,39-48^ consequently, O-GlcNAc is a potential mediator of metabolic memory.

We first demonstrated that the dnOGAh ON/OFF model as a novel approach for examining the potential role of cardiomyocyte O-GlcNAc in mediating metabolic memory (Figure 2A, 2B, S1A, S1B, S2A-S2D). Consistent with many other studies, we demonstrated that TAC resulted in increased cardiac hypertrophy, fibrosis and decreased systolic function (Figure 3).^37,57,58,86-89^ We found no significant increase in cardiac O-GlcNAc levels in response to TAC, which is in contrast to several other studies.^37,86,88,89^ However, other reports show no significant increase in O-GlcNAc following TAC,^87,88^ consistent with our own results (Figure 2A-2B). This variability in changes in O-GlcNAc levels in response to TAC is likely due to the variety of different pressure overload paradigms using TAC. For example, Zhu et. al. showed that cardiac O-GlcNAc increases at 1-wk. post-TAC but returned to control levels at 6-wks. post-TAC,^88^ pointing towards a relationship cardiac O-GlcNAc and the duration of TAC.

All measures of cardiac hypertrophy were exacerbated in the ON/OFF-TAC vs. Con-TAC groups(Figure 3A-3C and 3E), as defined by the magnitude of change between ON/OFF-TAC vs. ON/OFF-Sham groups is greater than that of the combination of main-effect changes either by Con-TAC vs. Con-Sham and/or ON/OFF-Sham vs. Con-Sham comparisons. We can also use basic synergy calculations to quantify exacerbation.^68^ Applying these criteria to our results, we see that in total heart weight, LV weight and LV cardiomyocyte cell area, the significant further increase between ON/OFF-TAC vs. ON/OFF-Sham groups is greater than the main-effect changes between the Con-TAC vs. Con-Sham and ON/OFF-Sham vs. Con-Sham groups. Exacerbation/synergy calculations for each show strong synergy for each measurement (*CI* = 0.66 for total heart weight, *CI* = 0.71 for LV weight, and *CI* = 0.63 for LV cell area). Transcriptomically, markers of cardiac hypertrophy also show exacerbated decrease in expression using our definition of exacerbation (Figure S8A, S8D-S8F). Cardiac fibrosis exacerbation is shown in perivascular fibrosis via histology (*CI* = 0.59, Figure 3D and 3F), and strengthened by exacerbated increase fibrosis marker gene expression (Figure S8B, S8G-S8I). With measures of cardiac dysfunction, we generally see exacerbated dysfunction; pulmonary edema showed exacerbated fluid backup into lungs (Figure 3G), and *Nppa* expression, as a classical marker of heart failure (HF), showed exacerbated increase in the ON/OFF-TAC vs Con-TAC (Figure S8C*, CI* = 0.36).

In summary, the phenotypic measurements of cardiac pathology in the current study support the concept that O-GlcNAc memory under subsequent TAC leads to exacerbated adverse LV cardiac hypertrophy, LV perivascular fibrosis, and non-systolic cardiac dysfunction; these results demonstrate the pathologic effects of “O-GlcNAc memory” at the cardiovascular pathology/systems level.

As discussed in the introduction, O-GlcNAc plays many roles in both transcriptional and epigenetic regulation;^31,41,42,44,45^ therefore, we performed mRNA sequencing of samples from the four groups of our study on O-GlcNAc memory with subsequent pressure-overload. We leveraged the scale of RNA sequencing data to start identifying the transcriptomic basis for potential driving molecular pathways of the exacerbated cardiac pathology seen in O-GlcNAc memory. Venn analysis of the DEGs filters and identifies DEGs changing in various overlapping combinations of the four key comparisons within O-GlcNAc memory with subsequent pressure-overload (Figure 4A). Venn diagram regions’ DEGs of interest were then imported into STRING-db to perform network and pathway analyses; the significantly enriched biologic pathways from this analysis, at a global level, are all related to the current study’s measures of exacerbated cardiac pathology such as cardiac hypertrophy, fibrosis and dysfunction (Figure 4B and 4C). Moreover, these pathways are potential underlying biological/molecular drivers of exacerbated cardiac pathology seen in O-GlcNAc memory with subsequent pressure-overload.

Investigating the DEGs of the enriched pathways, we identify specific DEGs which help place the general biological pathways in the specific context of cardiac pathology (Figure 5). The DEGs associated with ECM/fibrosis are genes encode proteins which can drive fibrotic remodeling (Figure 5A, 5B, S9A)—indicating potential molecular pathways behind the exacerbated fibrosis with known relevance to the heart.^90-94^ For example, *Ccn2* (connective tissue growth factor, CTGF, Figure 5A) is a known central mediator of pathologic fibrotic remodeling and can interact with TGFβ (with *Tgfb2* showing exacerbated increase expression in our data, Figure S9A) to induce and maintain fibrotic remodeling;^95,96^ and diabetic patients have 3x greater urine/plasma CTGF ratio compared to healthy subjects,^97^ supporting clinical relevance of CTGF in diabetic patients. Additionally, CTGF is known to stimulate adhesion, migration and growth of various cell types associated with cardiac remodeling, such as myofibroblasts, fibroblasts and smooth muscle cells;^98-100^ In the significantly enriched pathways for down-regulated genes, the majority of DEGs are associated with heart contraction/function and ion transport and show exacerbated downregulation (Figure 5C-5G, S9B and S9C). Specifically, many DEGs encode proteins for various cation channels and hormonal regulation of cardiac rhythm. In candidate DEGs associated with calcium signaling, many show exacerbated downregulation (Figure 5H, 5I, S9D). Importantly, the proteins encoded by these DEGs are known to be regulated (directly or indirectly) by O-GlcNAc and associated with the diabetic state—again demonstrating potential molecular pathways by which O-GlcNAc memory affects the cardiac environment.^39,101-106^

Furthermore, we also compared DEGs from our current study to the results of our previous work on sustained cardiomyocyte O-GlcNAcylation which identified and validated changes in redox mechanisms, OXPHOS function and mitochondrial homeostasis.^38^ In our current study, we see changes associated with an imbalance in the antioxidant vs ROS production. While *Nox4* expression is not significantly regulated by Con-TAC vs. Con-Sham and ON/OFF-Sham vs. Con-Sham, *Nox4* expression showed significantly increases in ON/OFF-TAC vs. ON/OFF-Sham groups (Figure 6A). NOX4 is a cardiac ROS producer implicated in both metabolic syndromes and pressure overload.^76-83^ Moreover, NOX4 is transcriptionally regulated,^81,107^ thus protein/activity can be directly inferred from the level of gene expression. *Cybb* (Figure S10A) encoding the NOX2 isoform is well studied in the context of hyperglycemia and O-GlcNAc, with work in Lu. et. al. (2020) showing regulation of NOX2 post-transcriptomically through O-GlcNAcylation of CAMKIIδ which activates NOX2 via phosphorylation.^103^ Conversely, the direct regulatory mechanisms of *Nox4* expression are still not very well understood; thus, with the literature surrounding NOX4 and it’s family members, and our results of exacerbation in the context of O-GlcNAc memory and TAC, strong opportunities exist within research into *Nox4/*NOX4 regulation. On the other side of redox mechanisms, *Sod2* (mitochondrial superoxide dismutase 2) shows exacerbated decrease in expression (Figure 6B). And *Nfe2l2* (NRF2), a central regulator of redox pathways and known O-GlcNAcylation target,^84^ is only upregulated in the Con-TAC vs. Con-Sham comparison (Figure 6C).

With regards to changes in cardiac metabolism, DEGs the current study associated with proteins of glucose metabolism show exacerbated downregulation (Figure 6D and 6E). Downstream, we investigated genes associated with enzymes of the citric acid cycle. *Cs*, encoding citrate synthase (CS), shows significantly exacerbated downregulation in expression (*CI* = 0.90, Figure 6F); and the majority of DEGs associated with TCA cycle enzymes show significantly exacerbated decrease in expression (Figure S11). Next, we investigated genes associated with proteins of the OXPHOS complexes. In the candidate DEG approach, we see exacerbated decrease in expression (Figure 6G-6K); and in the pathway-based approach, we see global decrease gene expression across all OXPHOS complexes in the ON/OFF-TAC vs. Con-TAC comparison (Figure S12). Finally, with all the changes in cardiac metabolism genes, *Ppargc1a* expression shows exacerbated decrease in gene expression (Figure S10C). Taking the results of our candidate DEG analysis, we show that changes in redox mechanisms (specifically in NOX4) could drive pathology exacerbation, with opportunities to further research into NOX4’s role in cardiac O-GlcNAcylation; additionally, exacerbated changes in gene expression associated with cardiac metabolism without compensation in mitochondrial homeostasis pathways could represent exacerbated changes due to O-GlcNAc memory with pressure-overload and/or drivers of exacerbated cardiac pathology.

With the novel results and strengths of our current study, one caveat that we have reiterated throughout the manuscript is that our initial foray into potential molecular pathways driving of O-GlcNAc memory are at the gene expression level; despite this caveat, it is encouraging to see the pathways and candidate DEGs from our RNA sequencing reinforce our phenotypic results of pathology exacerbation. Moving forward, the transcriptomic results set the initial blueprint and justification for future molecular and functional studies to elucidate the mechanisms underlying exacerbated cardiac pathology in transiently increased cardiomyocyte O-GlcNAc upon subsequent pressure overload. Additionally, with research showing epigenetic mechanisms potentially driving metabolic memory effects,^18,19^ future studies into DNA methylation are needed to investigate potential epigenetic mechanisms underlying O-GlcNAc memory. Beyond the transcriptomic limitations on mechanistic conclusions, other limitations exist within our current study; however, they also pose opportunities for future studies. Technically, due to limitations of our Vevo 770 system, we could not reliably obtain diastolic function measures; however, with our results on cardiac dysfunction, we recommend researchers studying O-GlcNAc memory evaluate diastolic function. Additionally, while the measures in our current study show exacerbated cardiac pathology, exercise tolerance testing is needed to further establish a more clinical vignette of CVD and failing hearts.

Nevertheless, the current study demonstrates previous transiently increased cardiomyocyte protein O-GlcNAcylation is sufficient to lead to exacerbation of cardiac pathology in context of pressure overload, and results support the concept of “O-GlcNAc memory” as a key player within the cardiac “metabolic memory” we see in diabetic patients. Our transcriptomic insights into the exacerbation begin progress within the pressing need for further specific research into the biologic mechanisms underpinning metabolic memory. This need reminds us of the goal of bringing our research to therapy; therefore, we need to continue to study pathophysiologic mechanisms and targets which can become future successful therapies to ameliorate the increased CVD associated with cardiac “metabolic memory”.

## Nonstandard Abbreviations and Acronyms

CVD: cardiovascular disease
DCCT: Diabetes Control and Complications Trial
DEG: differentially expressed genes
dnOGA: dominant-negative O-GlcNAcase
DOX: doxycycline
EDIC: Epidemiology of Diabetes Interventions and Complications
MACE: major adverse cardiovascular event
O-GlcNAc: O-linked β-N-acetylglucosamine
OGA: O-GlcNAcase
OGT: O-GlcNAc transferase
PTM: post-translational modification
TAC: transverse aortic constriction

## Acknowledgements

Dr. Michael Crowley of the UAB Heflin Center for Genomic Sciences, Genomics core laboratory. UAB Comparative Pathology Laboratory. Dr. Hind Lal lab for access to light microscope for histology slide image capture. Dr. Benjamin Lin for technical assistance with RNA sequencing analyses.

## Supplemental Materials

Data Supplement Tables S1–S7

Data Supplement Figures S1-S12

Data Supplement Raw Unedited Blots

Major Resources Table

ARRIVE Check List

## SUPPLEMENTAL MATERIALS

### DATA SUPPLEMENT TABLES

**Table S1.** Mouse survival analysis of all animals from current study.

Survival Analysis by Sex
| Sex | N | Deaths |
| --- | --- | --- |
| Female | 91 | 3 |
| Male | 84 | 7 |

Survival Analysis by Experimental Group
| Group | N | Deaths |
| --- | --- | --- |
| Con-Sham | 40 (F= 20, M=20) | 0 |
| ON/OFF-Sham | 31 (F=18, M=13) | 1 |
| Con-TAC | 52 (F= 27, M=25) | 6 |
| ON/OFF-TAC | 52 (F=26, M=26) | 3 |

Log-Rank (Mantel-Cox) Test: $X^2 = 6.116$ , $df = 3$ , $p\text{-value} = 0.1061$ , $p\text{-value summary} = NS$

**Table S2.** Other Echocardiography measures/parameters for Con-Sham, Con-TAC, ON/OFF-Sham and ON/OFF-TAC experimental groups at 8-weeks after TAC surgery, related to Figure S5.

| Experimental Group | Sex | Units Measure | mmHg. Peak Gradient | mm LVID;s | mm LVPW;s | mm IVS;s | mL/min CO |
| --- | --- | --- | --- | --- | --- | --- | --- |
| Con-Sham | Combined |  | 5.49 ± 0.53 | 2.36 ± 0.07 | 1.26 ± 0.04 | 1.28 ± 0.07 | 20.80 ± 1.30 |
|  | F |  | 5.86 ± 0.89 | 2.19 ± 0.08 | 1.23 ± 0.07 | 1.36 ± 0.08 | 20.93 ± 2.38 |
|  | M |  | 5.24 ± 0.69 | 2.52 ± 0.11 | 1.28 ± 0.05 | 1.21 ± 0.12 | 20.69 ± 1.95 |
| Con-TAC | Combined |  | 83.13 ± 2.66**** | 2.65 ± 0.09 | 1.42 ± 0.06 | 1.34 ± 0.10 | 15.51 ± 1.04* |
|  | F |  | 83.11 ± 4.32**** | 2.51 ± 0.10 | 1.51 ± 0.07 | 1.42 ± 0.12 | 14.54 ± 0.85 |
|  | M |  | 83.15 ± 3.61**** | 2.90 ± 0.14 | 1.27 ± 0.09 | 1.21 ± 0.17 | 17.18 ± 2.41* |
| ON/OFF-Sham | Combined |  | 6.53 ± 0.53 | 2.38 ± 0.09 | 1.29 ± 0.05 | 1.26 ± 0.09 | 20.58 ± 1.57 |
|  | F |  | 7.05 ± 0.71 | 2.26 ± 0.12 | 1.24 ± 0.07 | 1.29 ± 0.09 | 21.50 ± 2.31 |
|  | M |  | 6.18 ± 0.75 | 2.56 ± 0.12 | 1.36 ± 0.05 | 1.22 ± 0.17 | 19.20 ± 1.95 |
| ON/OFF-TAC | Combined |  | 83.19 ± 1.72**** | 2.70 ± 0.08 | 1.53 ± 0.06 = | 1.52 ± 0.09 | 14.87 ± 0.92* |
|  | F |  | 80.40 ± 3.00**** | 2.56 ± 0.09 | 1.52 ± 0.09 | 1.56 ± 0.12 | 13.69 ± 1.14 |
|  | M |  | 85.97 ± 1.13**** | 2.91 ± 0.10 | 1.54 ± 0.05 * | 1.46 ± 0.17 | 16.98 ± 17.00* |

Left Ventricle Internal Dimension at Systole (LVID;s), Left Ventricle Posterior Wall Thickness at Systole (LVPW;s), Interventricular Septum Thickness at Systole (IVS;s), Cardiac Output (CO). Data presented as Mean ± SEM. Statistical significance was based on two-way ANOVA followed by Tukey's multiple comparisons test (\* $p < 0.05$ , \*\*\*\* $p < 0.0001$ for -TAC vs. -Sham Control; = $p < 0.05$ for ON/OFF-TAC vs. Con-TAC).

Table S3. All differentially expressed genes for each pairwise comparison between experimental groups.

Table S4: mRNA sequencing sample ID and conditions/groups

**Table S5.**
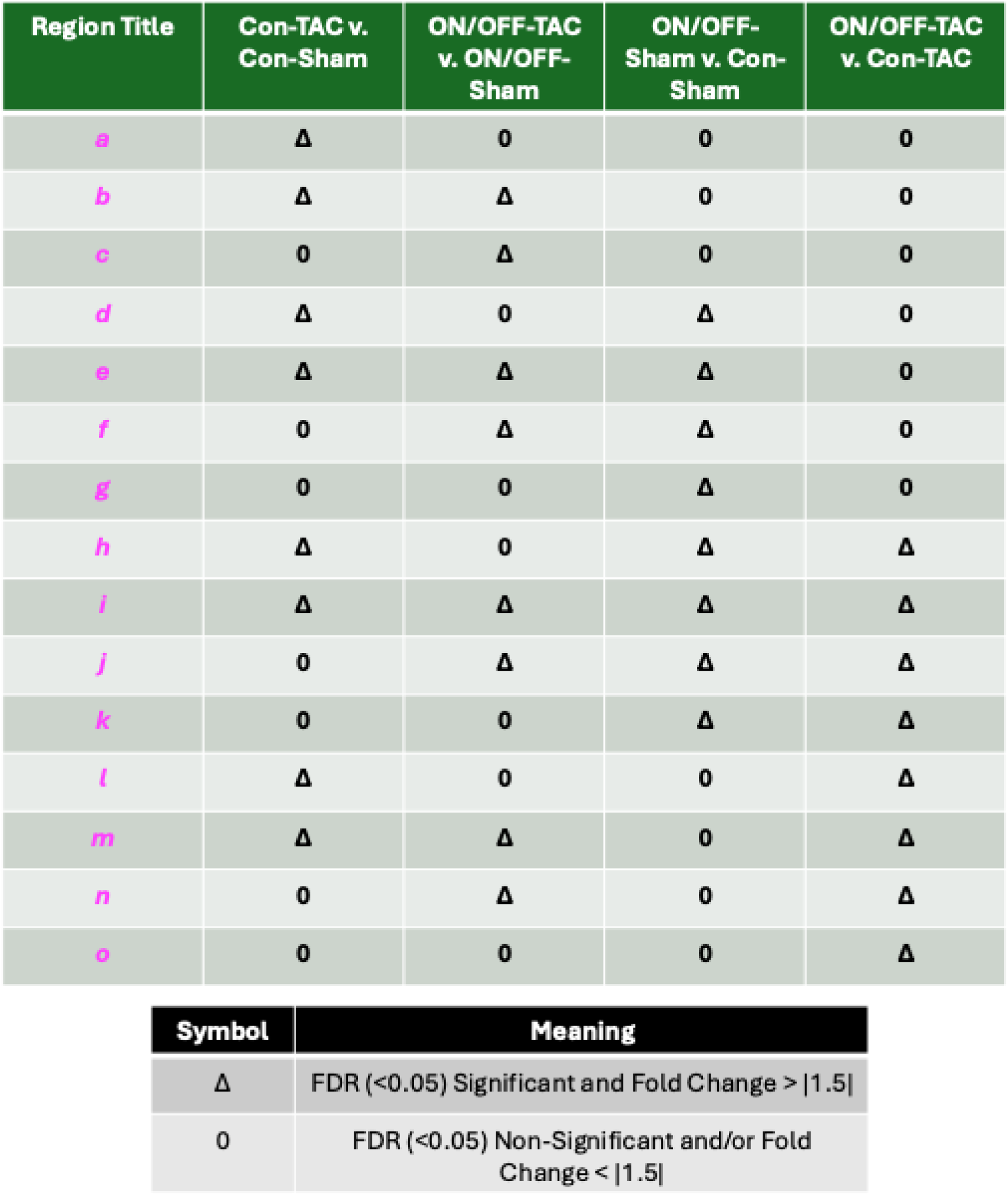
Key of 4-way Venn diagram comparison regions.

Table S6: All DEGs from each region of the 4-way Venn analysis.

Table S7: Tables of all significantly enriched pathways from the 4-way Venn regional DEGs using STRING-db.

### DATA SUPPLEMENT FIGURES

**Figure S1, related to Figure 2.**
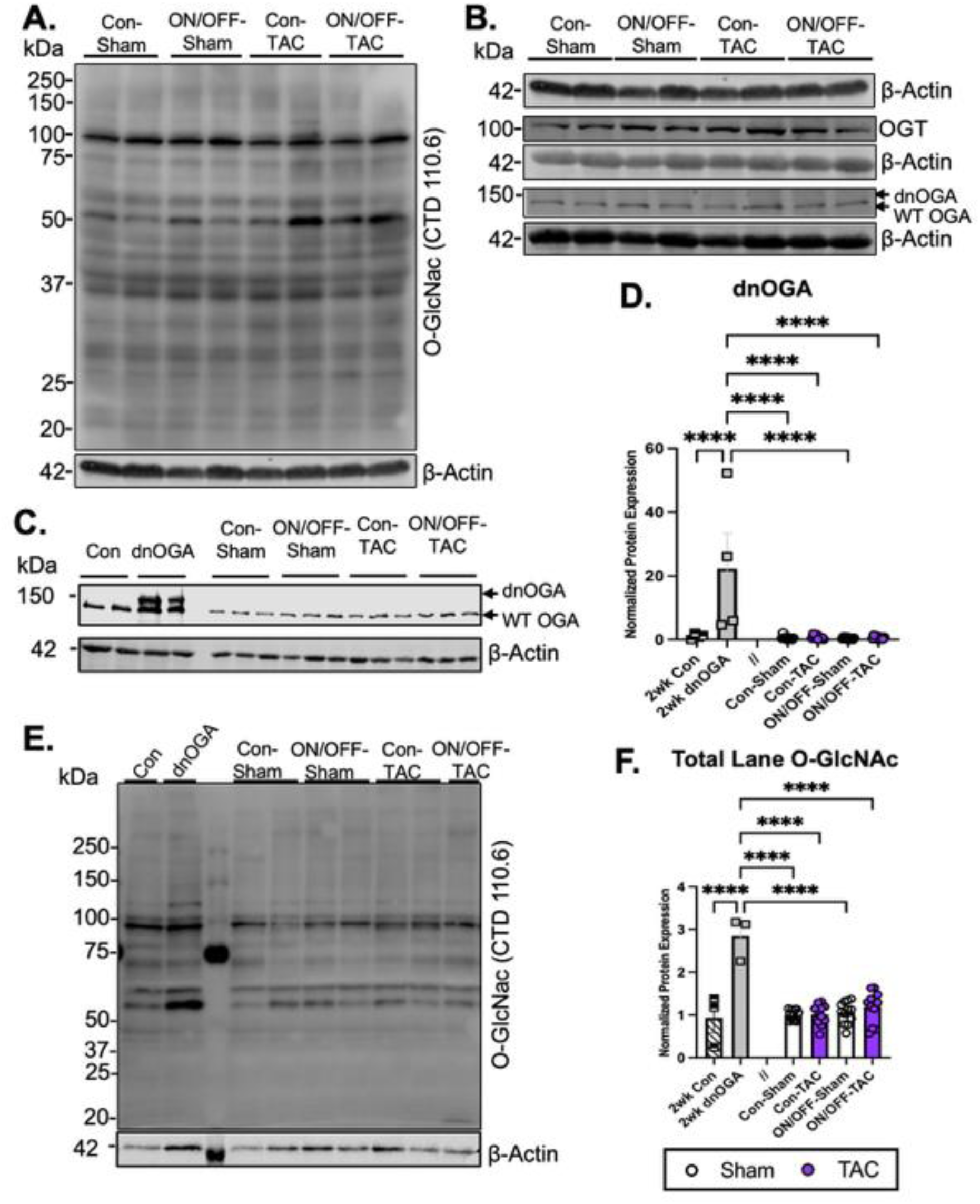
dnOGAh ON/OFF model confirmation: dnOGA transgene and O-GlcNAc quantification. **A-B)** Un-arranged Western blot membranes image from Figure 2A. **C-D)** Western blotting of left ventricular (LV) whole cell lysate for endogenous/WT OGA and dnOGA with densitometry quantification in 2-wk DOX treated Control, 2-wk DOX induced dnOGA, and four groups from the dnOGAh ON/OFF-TAC paradigm (Con-Sham, ON/OFF-Sham, Con-TAC, ON/OFF-TAC). **E-F)** Western blot assay of left ventricle (LV) whole cell protein extract for protein O-GlcNAcylation (via CTD110.6) with densitometry quantification in 2-wk DOX treated Control, 2-wk DOX induced dnOGA, and four groups from the dnOGAh ON/OFF-TAC paradigm (Con-Sham, ON/OFF-Sham, Con-TAC, ON/OFF-TAC). All quantitative data are mean ± S.E.M; Statistical significance was based on two-way ANOVA followed by Tukey’s multiple comparisons test (**** *p* < 0.0001).

**Figure S2. related to Figure 2.**
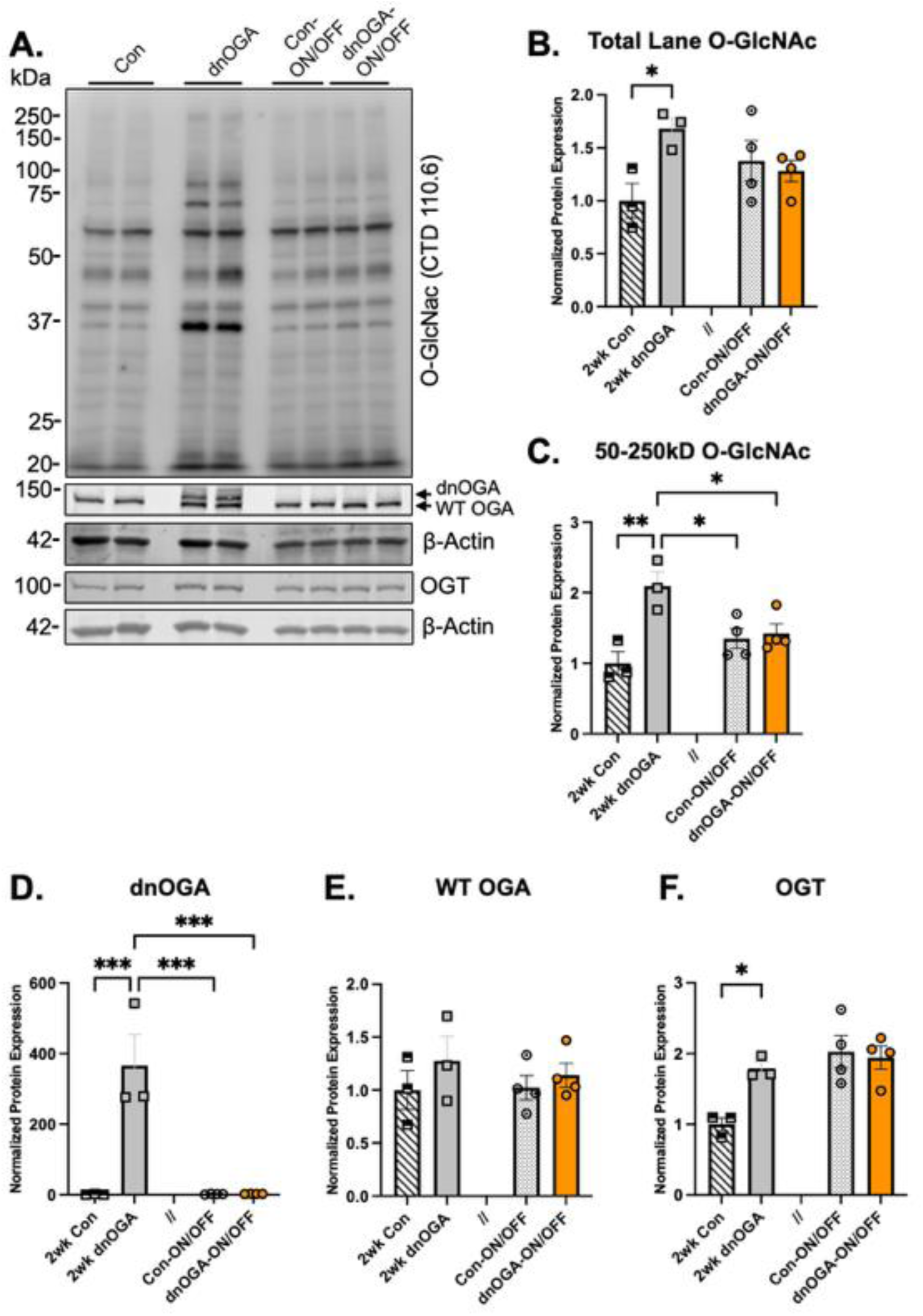
dnOGAh ON/OFF model confirmation: dnOGA transgene and O-GlcNAc quantification after “ON/OFF”. **A)** Western blotting of left ventricular (LV) whole cell lysate for: protein O-GlcNAcylation (via CTD110.6), endogenous/WT OGA and dnOGA, and OGT with densitometry quantification in 2-wk DOX treated Control, 2-wk DOX induced dnOGA (for “ON” branch), 2-wk DOX then 2-wk standard chow treated Control (Con-ON/OFF), and 2-wk DOX then 2-wk standard chow induced dnOGA (dnOGA-ON/OFF) conditions. **B-F)** Densitometry analysis of all immunoblots. **B)** Quantification of whole-lane cardiac protein O-GlcNAcylation (20–250 kDa protein size). **C)** Quantification of cardiac protein O-GlcNAcylation at 50–250 kDa protein size. **D**) Quantification of dnOGA transgene expression. **E)** Quantification of endogenous/WT OGA protein. **F)** Quantification of OGT protein. All quantitative data are mean ± S.E.M; Statistical significance was based on two-way ANOVA followed by Tukey’s multiple comparisons test (* *p* < 0.05, ** *p* < 0.01, *** *p* < 0.001).

**Figure S3.**
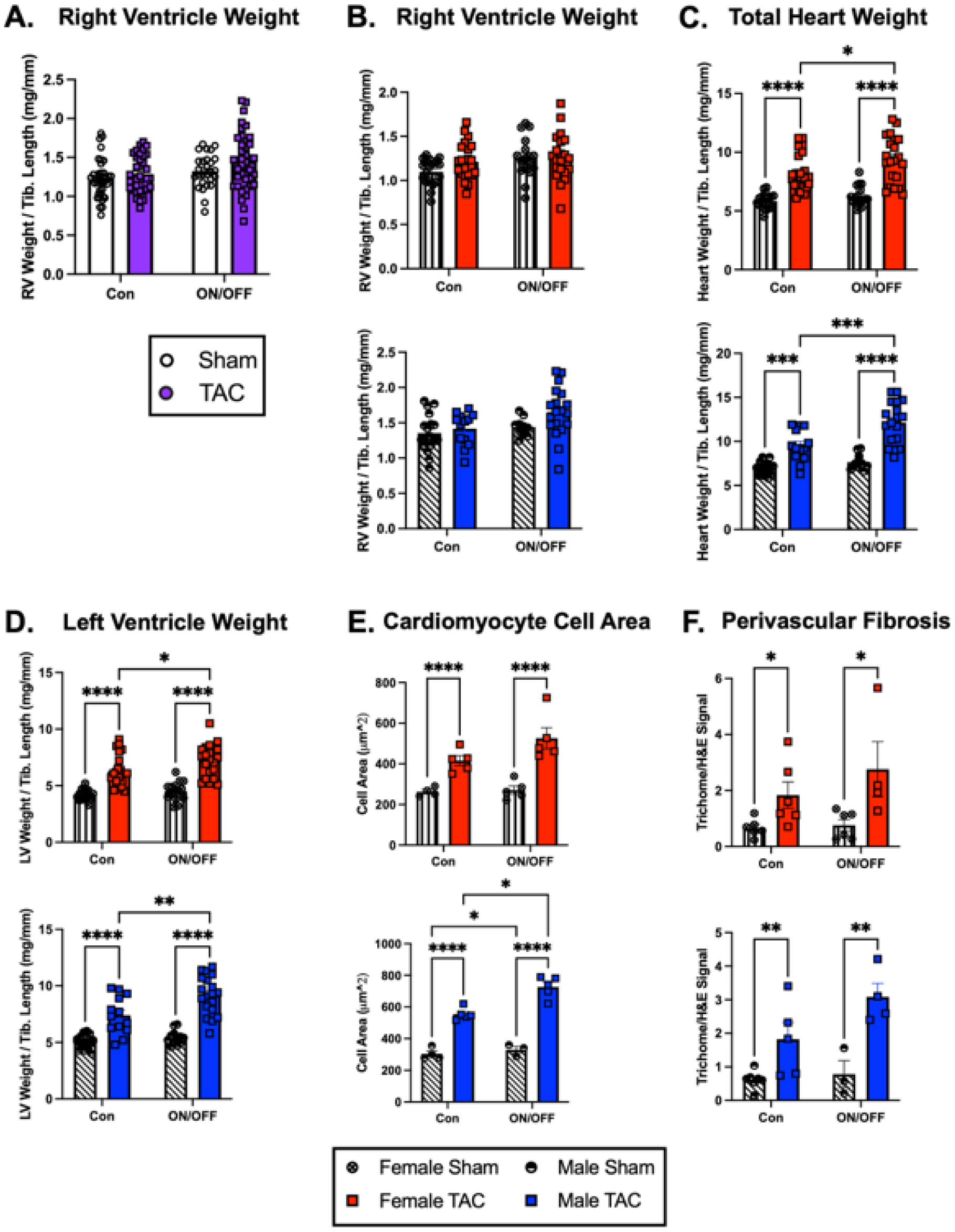
Transient Cardiac O-GlcNAc induction with subsequent pressure-overload induces cardiac hypertrophy and fibrosis by sex. **A)** Normalized ratio of right ventricle (RV) to tibia length for Con-TAC and ON/OFF-TAC mice compared to age-matched Sham controls (Con-Sham and ON/OFF-Sham) (*n* ≥ 27). **B-D**) Normalized ratio of total heart weight to tibia length (TL) (**B**), left ventricle (LV) plus septum to TL (**C**), and right ventricle (RV) to TL (**D**), for Con-TAC and ON/OFF-TAC mice compared to age-matched Sham controls (Con-Sham and ON/OFF-Sham) split by sex. **E)** Average cross-sectional LV cardiomyocyte cell area for Con-TAC and ON/OFF-TAC mice compared to age-matched Sham controls (Con-Sham and ON/OFF-Sham) split by sex. **F)** Normalized ratio of trichrome to H&E signal of perivascular LV tissue for Con-TAC and ON/OFF-TAC mice compared to age-matched Sham controls (Con-Sham and ON/OFF-Sham) split buy sex. Statistical significance was based on two-way ANOVA followed by Tukey’s multiple comparisons test (* *p* < 0.05, ** *p* < 0.01, *** *p* < 0.001, **** *p* < 0.0001), and unpaired non-parametric Mann-Whitney (two tailed) (# *p* < 0.05). All calculated and analysis data are mean ± S.E.M.

**Figure S4, related to Figure 3.**
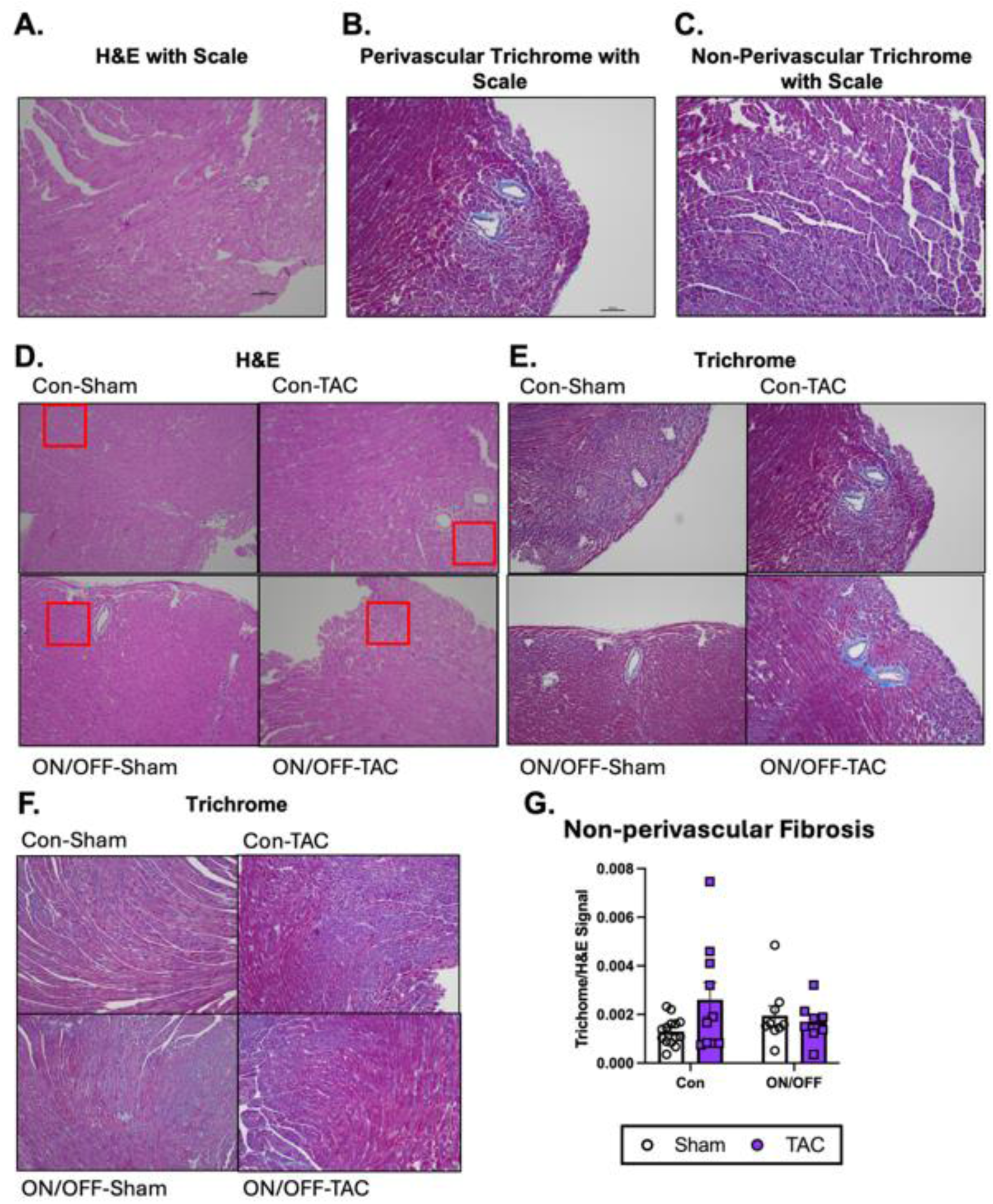
Additional LV fibrosis and cellular hypertrophy measures and controls. **A-C)** Representative images with scale bar for histological examination of LV cardiac tissue with H&E staining, perivascular LV cardiac tissue with H&E and trichrome staining, and interstitial LV cardiac tissue with H&E and trichrome staining. **D)** Uncropped/full images from representative images in Figure 3D for histological examination of LV cardiac tissue with H&E staining, with zoomed region of interest for cross-sectional cardiomyocyte cell area analysis frame in red. **E)** Uncropped/full images from representative images in Figure 4A for histological examination of perivascular LV cardiac tissue with H&E and trichrome staining. **F)** Representative images for histological examination of interstitial LV cardiac tissue with H&E and trichrome staining. **G)** Normalized ratio of trichrome to H&E signal of interstitial LV tissue for Con-TAC and ON/OFF-TAC mice compared to age-matched Sham controls (Con-Sham and ON/OFF-Sham) (*n* ≥ 8). Statistical significance was based on two-way ANOVA followed by Tukey’s multiple comparisons test. All calculated and analysis data are mean ± S.E.M.

**Figure S5.**
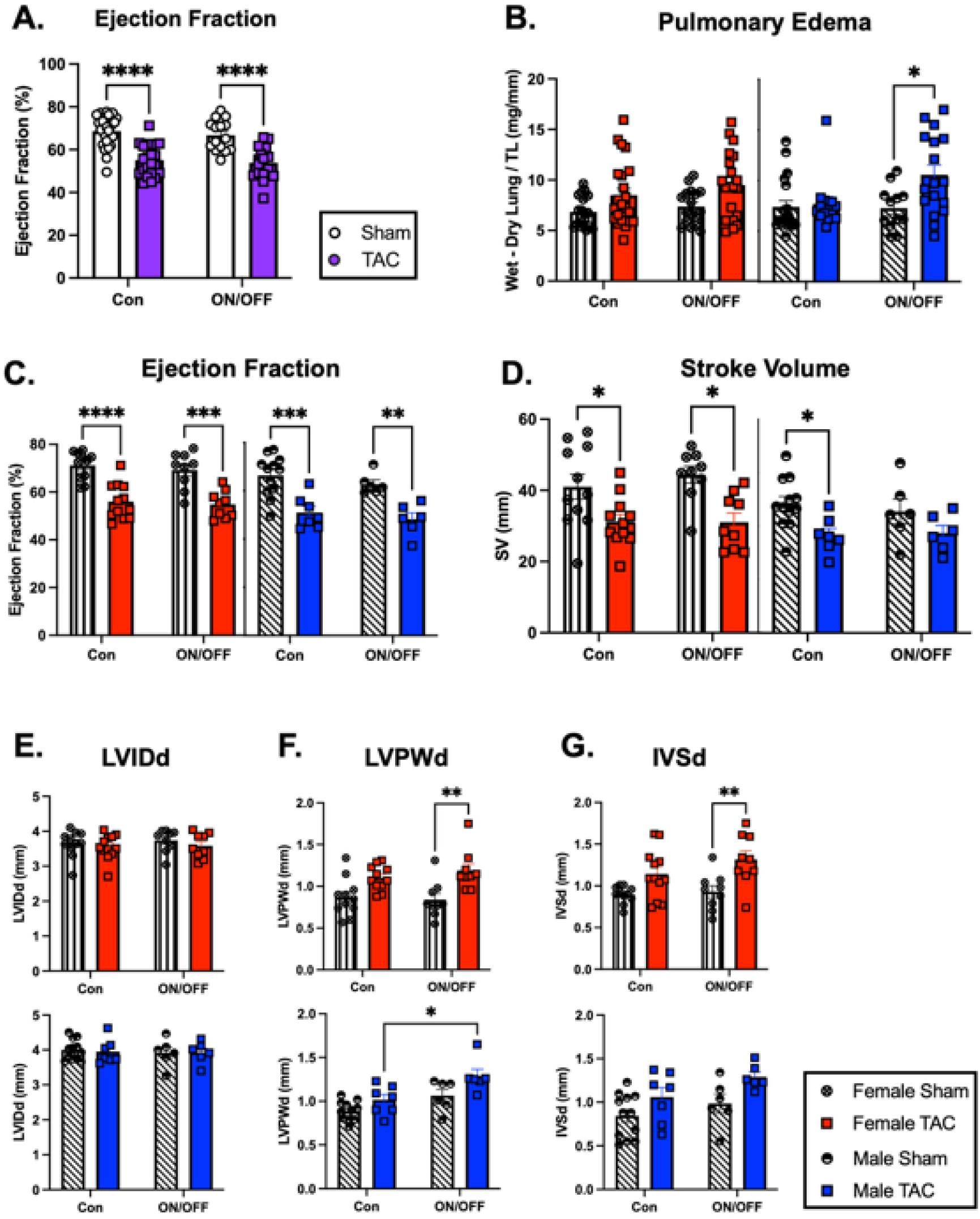
Transient Cardiac O-GlcNAc induction with subsequent pressure-overload associated with various measures of cardiac dysfunction by sex. **A)** Normalized ratio of wet-dry lung weight to TL for Con-TAC and ON/OFF-TAC mice compared to age-matched Sham controls (Con-Sham and ON/OFF-Sham) (*n* ≥ 29). **B)** Normalized ratio of wet-dry lung weight to TL for Con-TAC and ON/OFF-TAC mice compared to age-matched Sham controls (Con-Sham and ON/OFF-Sham) split by sex. **C-G)** Cardiac function measures via echocardiography split by sex for: cardiac systolic function as measured by LV ejection fraction from M-mode LV traces (**C**), stroke volume from M-mode LV traces (**D**), left ventricular internal dimension at diastole (LVIDd) (**E**), cardiac wall thickening as measured by LV posterior wall thickness at diastole (LVPWd) (**F**), and inter-ventricular septum thickness at diastole (IVSd) (**G**). Statistical significance was based on two-way ANOVA followed by Tukey’s multiple comparisons test (* *p* < 0.05, ** *p* < 0.01, *** *p* < 0.005). All calculated and analysis data are mean ± S.E.M.

**Figure S6.**
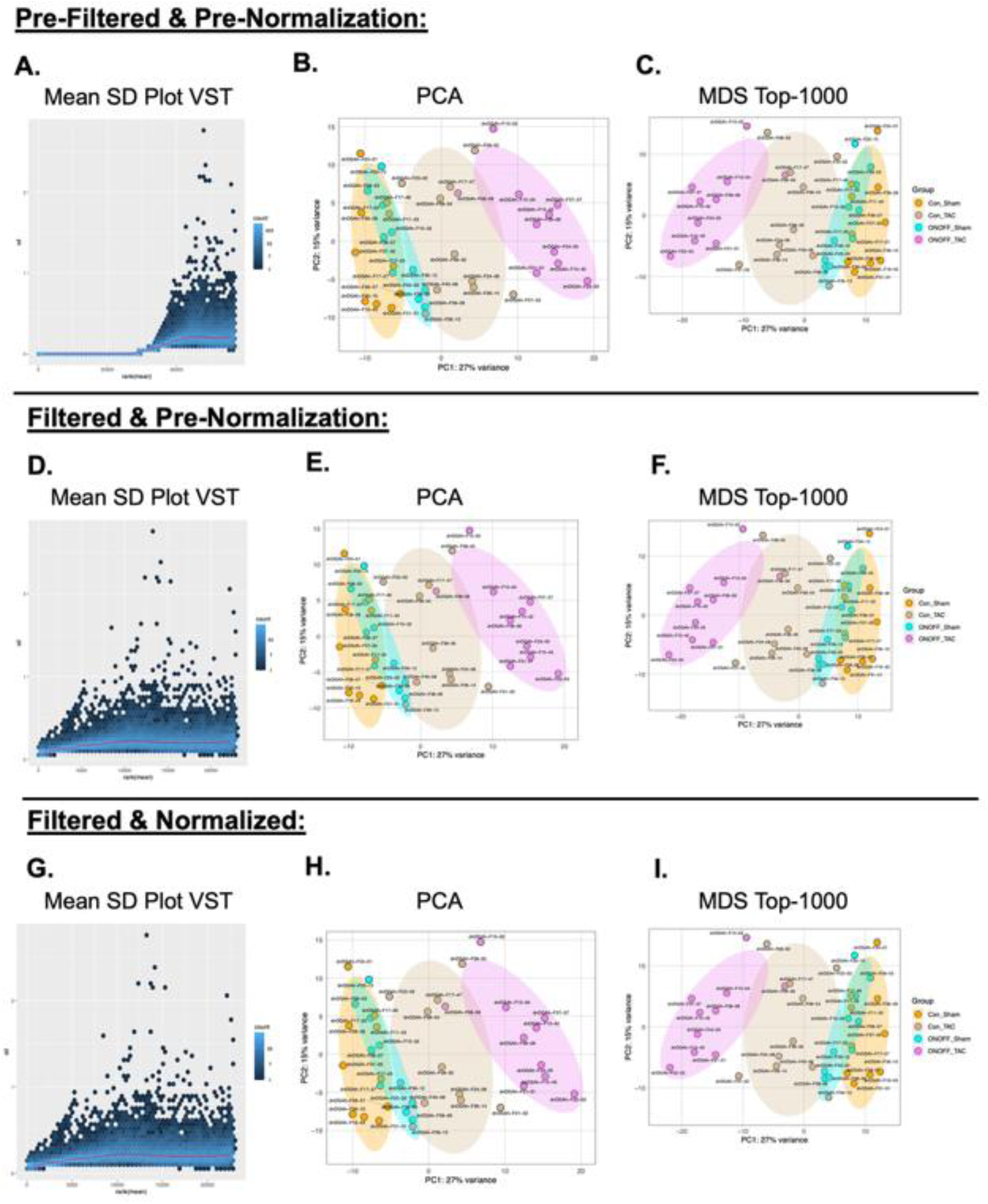
RNA Sequencing quality control. **A-C**) For pre-filtered and pre-normalized data from mRNA sequencing: **A)** Mean-standard deviation plot after variance stabilizing transformation to visualize level of homoscedasticity of the pre-filtered and pre-normalized data. **B)** Plot of unsupervised principal component analysis (PCA) utilizing regularized logarithm-transformed pre-filtered and pre-normalized read count data of all four conditions with labeled samples (Con-Sham: orange, Con-Tac: tan, ON/OFF-Sham: light blue, ON/OFF-TAC: pink). The plot denotes variance explained by first two principal components, PC1 and PC2. **C)** Plot of multidimensional scaling (MDS) with pre-filtered and pre-normalized read count data of all four conditions with labeled samples (Con-Sham: orange, Con-Tac: tan, ON/OFF-Sham: light blue, ON/OFF-TAC: pink). The plot denotes variance explained by first two principal components, PC1 and PC2 for MDS. **D-F**) For filtered and pre-normalized data from mRNA sequencing: **D)** Mean-standard deviation plot after variance stabilizing transformation to visualize level of homoscedasticity of the filtered and pre-normalized data. **E)** PCA plot utilizing regularized logarithm-transformed filtered and pre-normalized read count data of all four conditions with labeled samples (Con-Sham: orange, Con-Tac: tan, ON/OFF-Sham: light blue, ON/OFF-TAC: pink). The plot denotes variance explained by first two principal components, PC1 and PC2. **F)** MDS plot with filtered and pre-normalized read count data of all four conditions with labeled samples (Con-Sham: orange, Con-Tac: tan, ON/OFF-Sham: light blue, ON/OFF-TAC: pink). The plot denotes variance explained by first two principal components, PC1 and PC2 for MDS. **G-I**) For filtered and normalized data from mRNA sequencing: **G)** Mean-standard deviation plot after variance stabilizing transformation to visualize level of homoscedasticity of the filtered and normalized data. **H)** PCA plot utilizing regularized logarithm-transformed filtered and normalized read count data of all four conditions with labeled samples (Con-Sham: orange, Con-Tac: tan, ON/OFF-Sham: light blue, ON/OFF-TAC: pink). The plot denotes variance explained by first two principal components, PC1 and PC2. **I)** MDS plot with filtered and normalized read count data of all four conditions with labeled samples (Con-Sham: orange, Con-Tac: tan, ON/OFF-Sham: light blue, ON/OFF-TAC: pink). The plot denotes variance explained by first two principal components, PC1 and PC2 for MDS.

**Figure S7.**
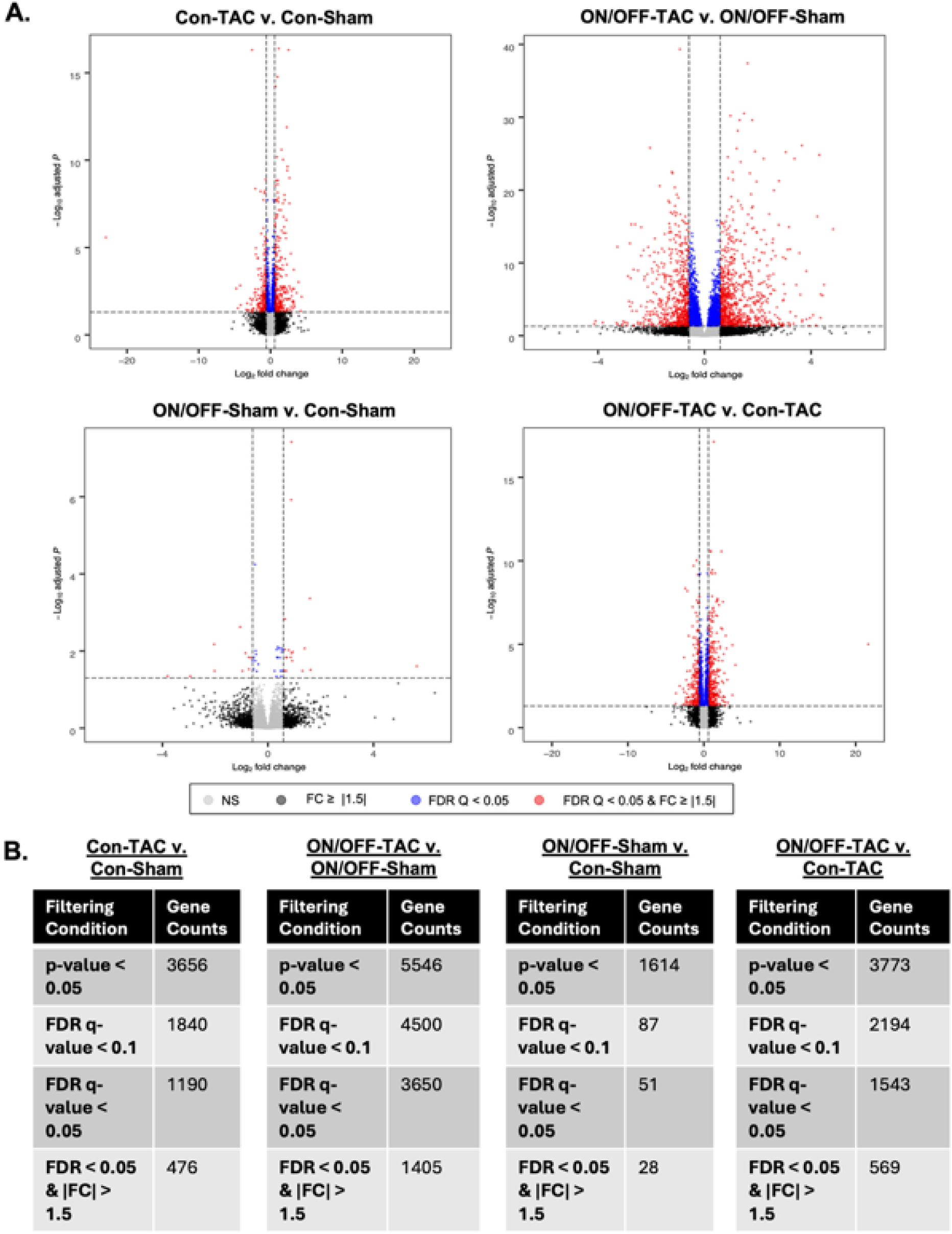
General transcriptome characterization (volcano plots, DEG frequency tables). **A)** Visual representation of DEGs via volcano plot of all genes with cutoffs at FDR/q < 0.05 and FC ≥ |1.5| for all four relevant pairwise comparisons between experimental groups (Con-TAC vs. Con-Sham, ON/OFF-TAC vs. ON/OFF-Sham, ON/OFF-Sham vs. Con-Sham, ON/OFF-TAC vs. Con-TAC). **B)** Frequency tables of DEGs at various significance cutoffs each of the relevant pairwise comparisons between experimental groups (Con-TAC vs. Con-Sham, ON/OFF-TAC vs. ON/OFF-Sham, ON/OFF-Sham vs. Con-Sham, ON/OFF-TAC vs. Con-TAC).

**Figure S8.**
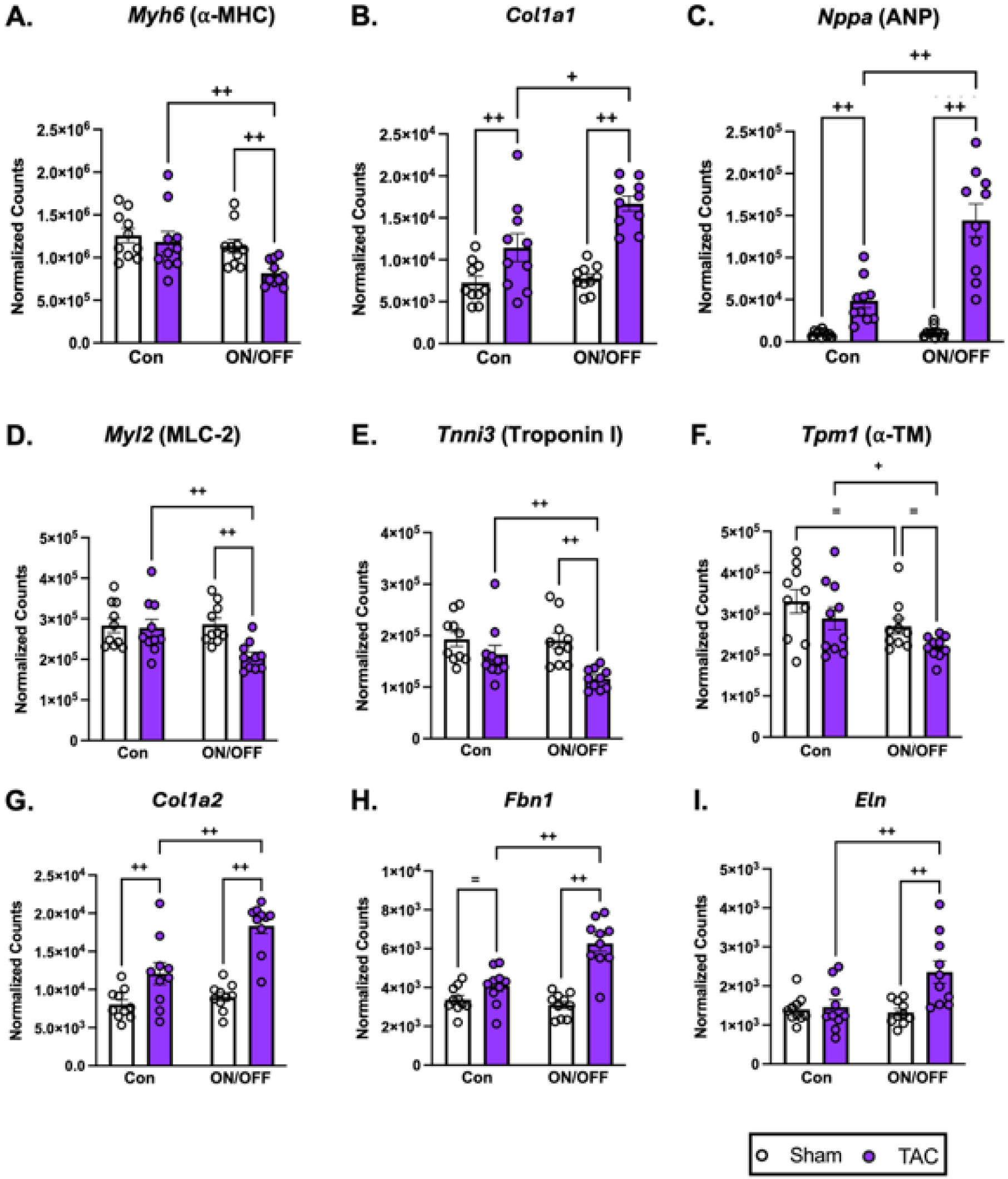
Extended candidate DEGs that reinforce phenotypic findings of pathology exacerbation. **A-I)** Specific candidate DEGs from mRNA sequencing of LV tissue (*n* = 5 per sex, per group) in normalized counts. **A)** myosin heavy chain 6 (*Myh6*). **B)** Collagen type 1 alpha-1 (*Col1a1).* **C)** Natriuretic peptide A (*Nppa*; ANP). **D)** myosin regulatory light chain 2 (*Myl2*). **E)** cardiac troponin I (*Tnni3*). **F)** tropomyosin α-1 chain (*Tpm1*). **G)** Collagen type 1 alpha-2 (*Col1a2).* **H)** fibrilin-1 (*Fbn1*). **I)** elastin (*Eln*). Statistical significance was based on Wald test (*= p* < 0.05, + *q* < 0.1, and ++ *q* < 0.05). All calculated and analysis data are mean ± S.E.M.

**Figure S9, related to Figure 5.**
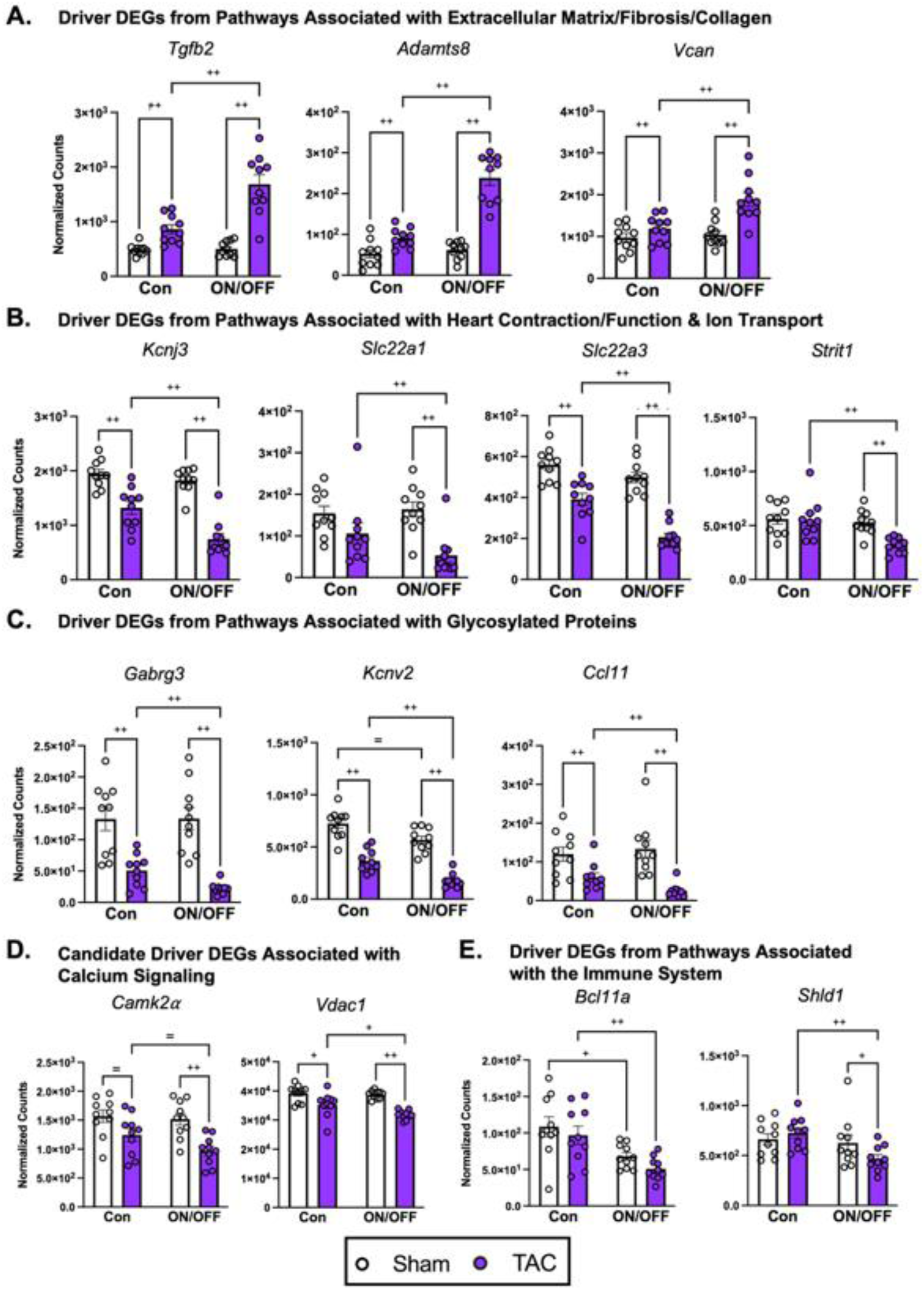
Extended DEGs from pathway analyses of key subsets and candidate DEGs that identify potential transcriptomic drivers of pathology exacerbation. **A-E)** Specific DEGs from mRNA sequencing of LV tissue (*n* = 5 per sex, per group) in normalized counts that were significantly enriched in pathway analysis of Figure 5. **A**) Specific DEGs from pathways associated with extracellular matrix/fibrosis/collagen: transforming growth factor-β2 (*Tgfb2*), ADAM metallopeptidase with thrombospondin type 1 motif 8 (*Adamts8*), and Versican (*Vcan*). **B**) Specific DEGs from pathways associated with heart contraction/function and ion transport: potassium channel (*Kcnj3*), organic cation transporters (*Slc22a1* and *Slc22a3*), and small transmembrane regulator of ion transport 1 (*Strit1*). **C**) Specific DEGs from pathways associated with glycosylated proteins: GABA type A receptor (*Gabrg3*), voltage-gate potassium channel subunit (*Kcnv2*), and Eotaxin-1 (*Ccl11*). **D**) Specific DEGs associated with calcium signaling: Calcium/calmodulin-dependent protein kinase type II subunit alpha (*Camk2α*), and Voltage-dependent anion channel 1 (*Vdac1*). **E**) Specific DEGs associated with the immune system: B-cell lymphoma/leukemia 11a (*Bcl11a*), Shieldin complex subunit 1 (*Shld1*). Statistical significance was based on Wald test (*= p* < 0.05, + *q* < 0.1, and ++ *q* < 0.05). All calculated and analysis data are mean ± S.E.M.

**Figure S10, related to Figure 6.**
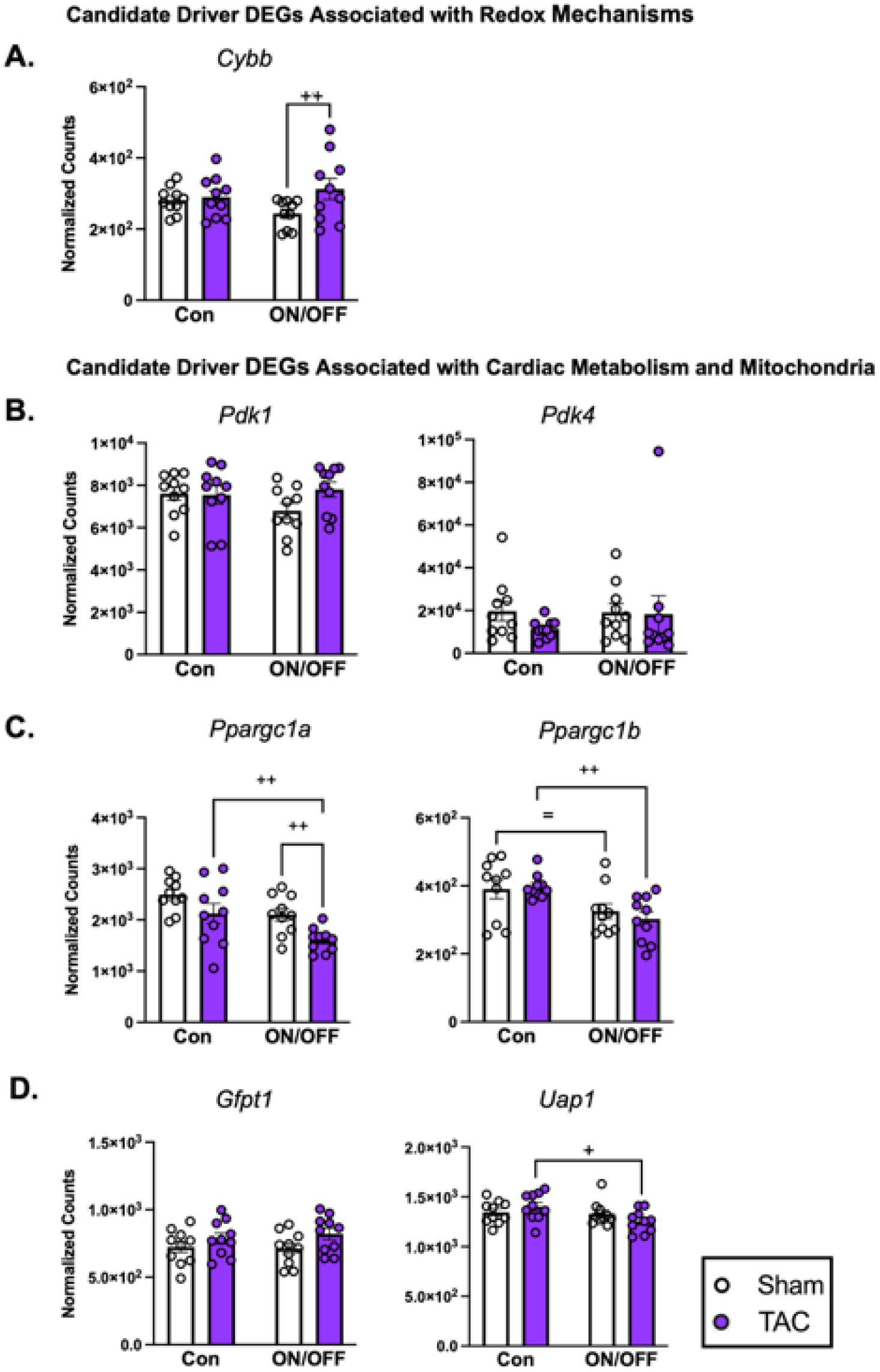
Extended DEGs from redox, metabolism, mitochondria related genes. **A-D)** Specific candidate DEGs from mRNA sequencing of LV tissue (*n* = 5 per sex, per group) in normalized counts. **A**) Specific DEGs associated with redox mechanisms: Cytochrome b beta (NOX2) (*Cybb*). **B**) Specific DEGs associated with cardiac metabolism: Pyruvate dehydrogenase kinase 1 (*Pdk1*), Pyruvate dehydrogenase kinase 4 (*Pdk4*). **C**) Specific DEGs associated with mitochondrial biogenesis/homeostasis: PPARG coactivator 1 alpha (*Ppargc1a*), PPARG coactivator 1 beta (*Ppargc1b*). **D**) Specific DEGs associated with the Hexosamine Biosynthesis Pathway (HBP): Glutamine-fructose-6-phosphate transaminase 1 (*Gfpt1*), UDP-N-acetylglucosamine pyrophosphorylase (*Uap1)*. Statistical significance was based on Wald test (*= p* < 0.05, + *q* < 0.1, and ++ *q* < 0.05). All calculated and analysis data are mean ± S.E.M.

**Figure S11, related to Figure 6.**
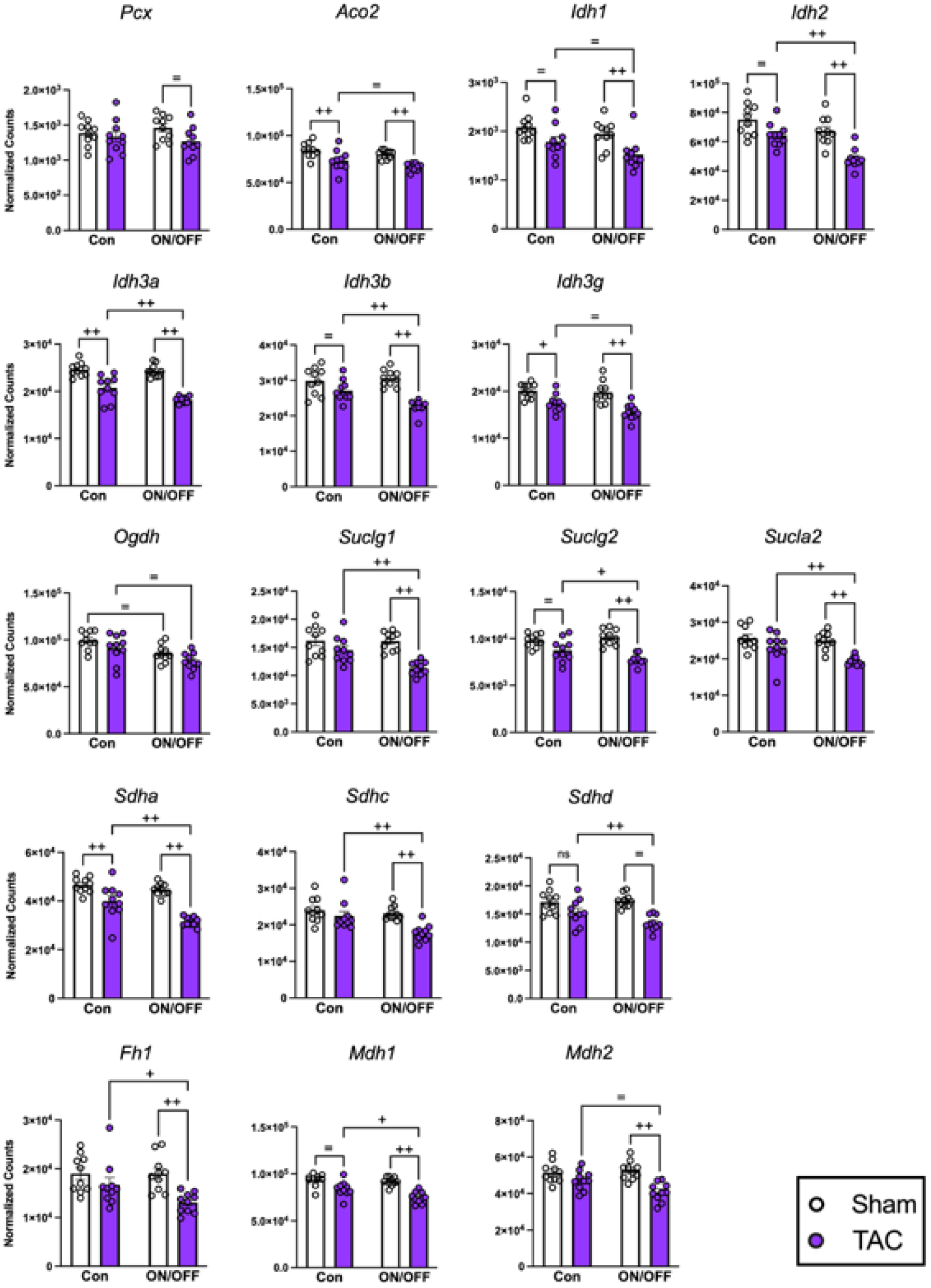
Additional DEGs from tricarboxylic acid cycle (TCA) related genes. Specific candidate DEGs from mRNA sequencing of LV tissue (*n* = 5 per sex, per group) in normalized counts. **A**) Specific DEGs associated with proteins/enzymes of the tricarboxylic acid (TCA) cycle: Pyruvate carboxylase (*Pcx*), Aconitase 2 (*Aco2*), Isocitrate dehydrogenase 1 (*Idh1*), Isocitrate dehydrogenase 2 (*Idh2*), Isocitrate dehydrogenase 3a (*Idh3a*), Isocitrate dehydrogenase 3b (*Idh3b*), Isocitrate dehydrogenase 3g (*Idh3g*), Oxoglutarate dehydrogenase (*Ogdh*), Succinate-CoA ligase alpha subunit (*Suclg1*), Succinate-CoA ligase GDP-forming beta subunit (*Suclg2*), Succinate-CoA ligase beta subunit (*Sucla2*), Succinate dehydrogenase subunit a (*Sdha*), Succinate dehydrogenase subunit c (*Sdhc*), Succinate dehydrogenase subunit d (*Sdhd*), Fumarate hydratase 1 (*Fh1*), Malate dehydrogenase 1 (*Mdh1*), Malate dehydrogenase 2 (*Mdh2*). Statistical significance was based on Wald test (*= p* < 0.05, + *q* < 0.1, and ++ *q* < 0.05). All calculated and analysis data are mean ± S.E.M.

**Figure S12, related to Figure 6.**
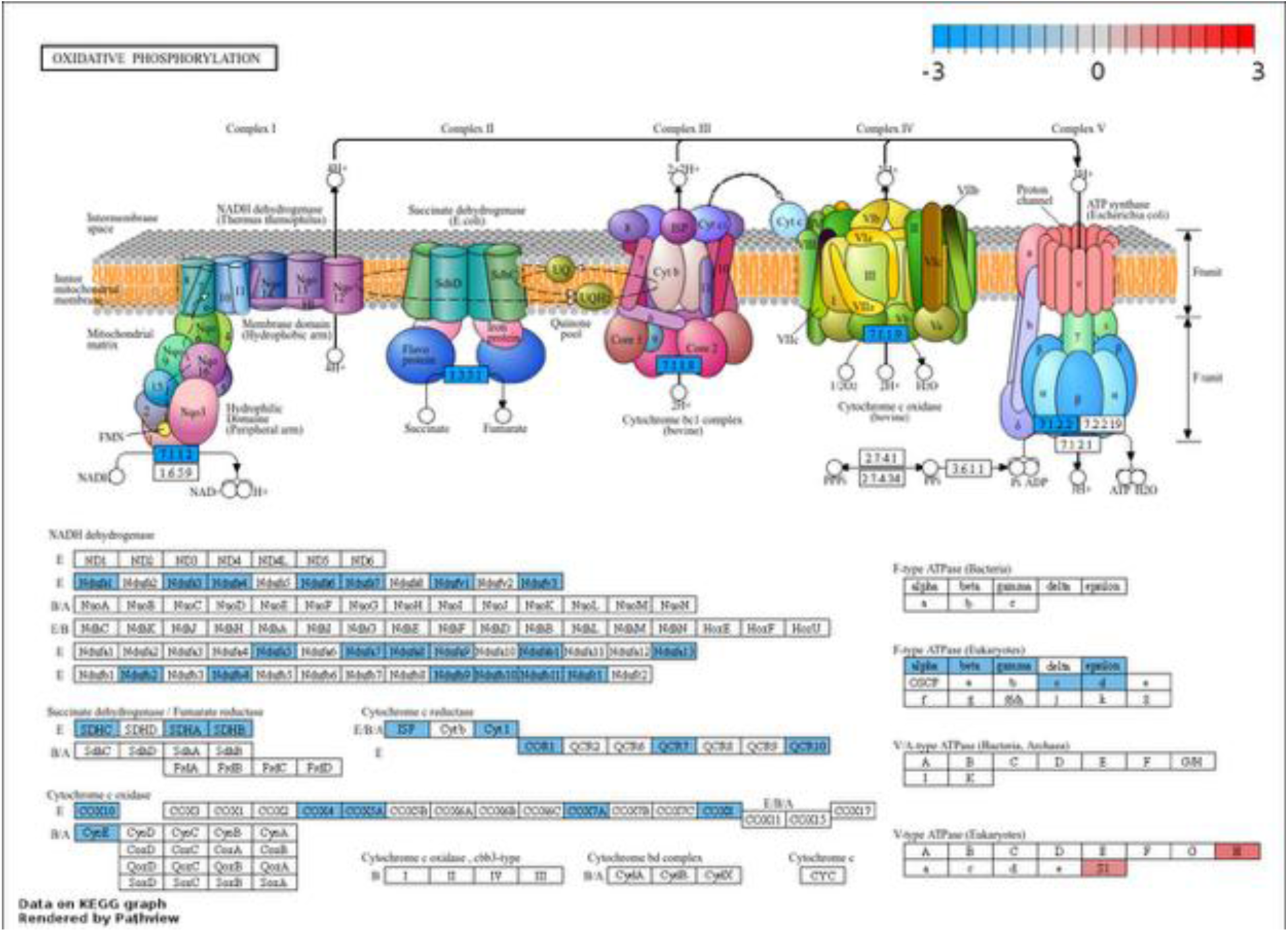
KEGG Pathview analyses of significant ON/OFF-TAC vs. Con-TAC DEGs shows overall decreased expression of mitochondrial electron transport chain genes. KEGG graph rendered by Pathview for significantly enriched KEGG pathway (Oxidative Phosphorylation) from import of significant (FDR/q < 0.05) DEGs from ON/OFF-TAC vs. Con-TAC comparison.

### DATA SUPPLEMENT RAW UNEDITED BLOTS AND HISTOLOGY IMAGES

**Figure 2, S1, and S2**

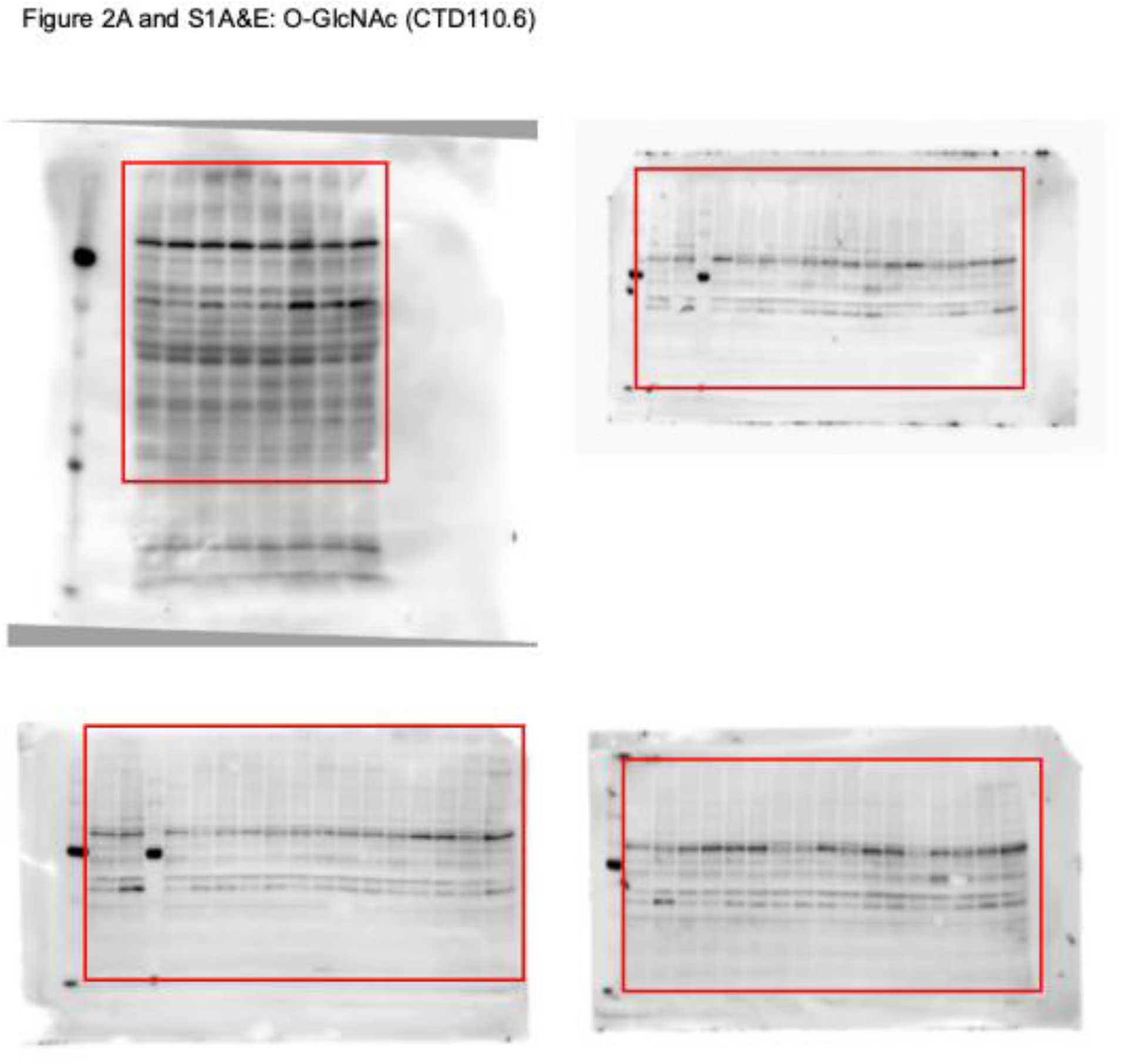

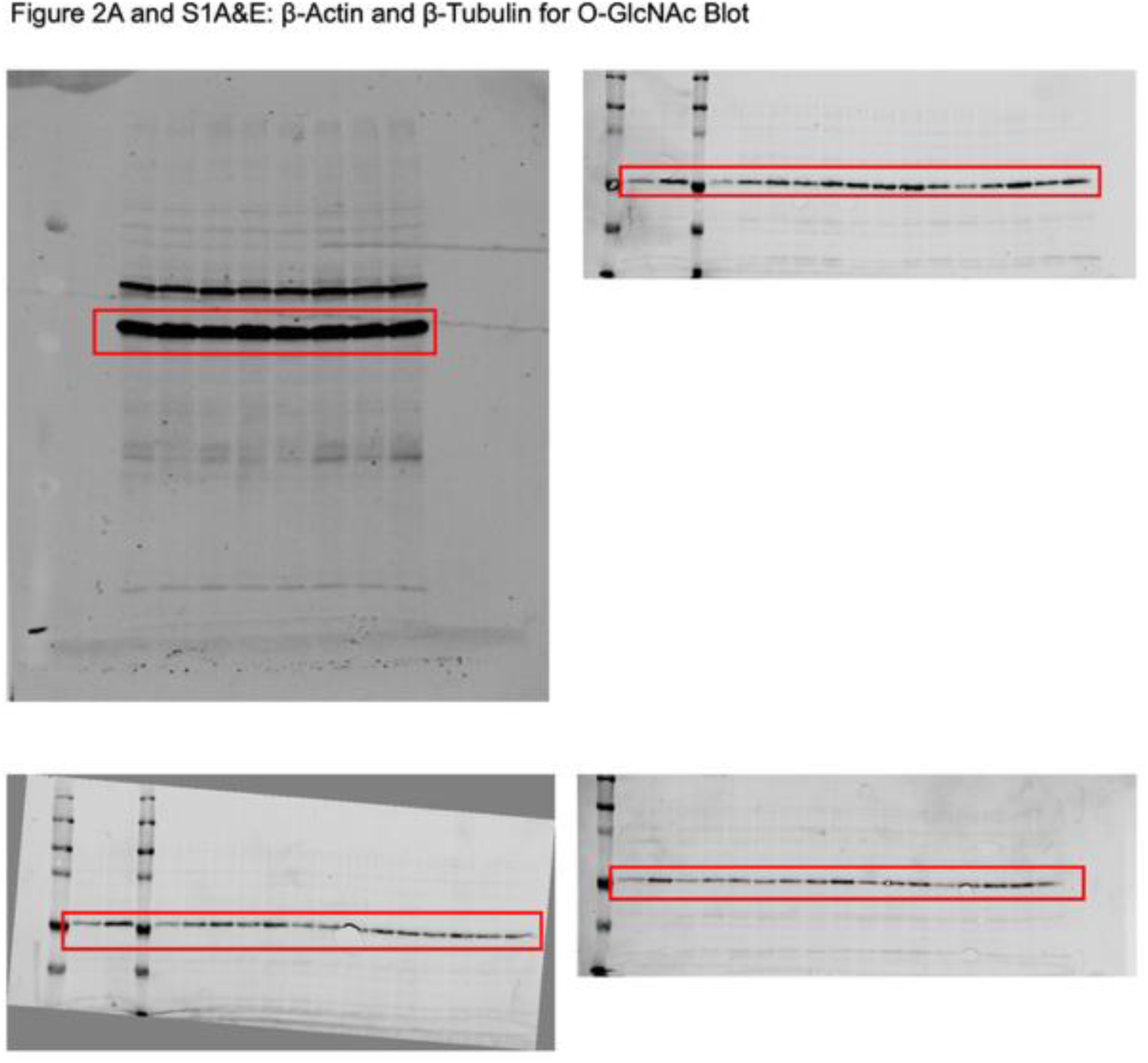

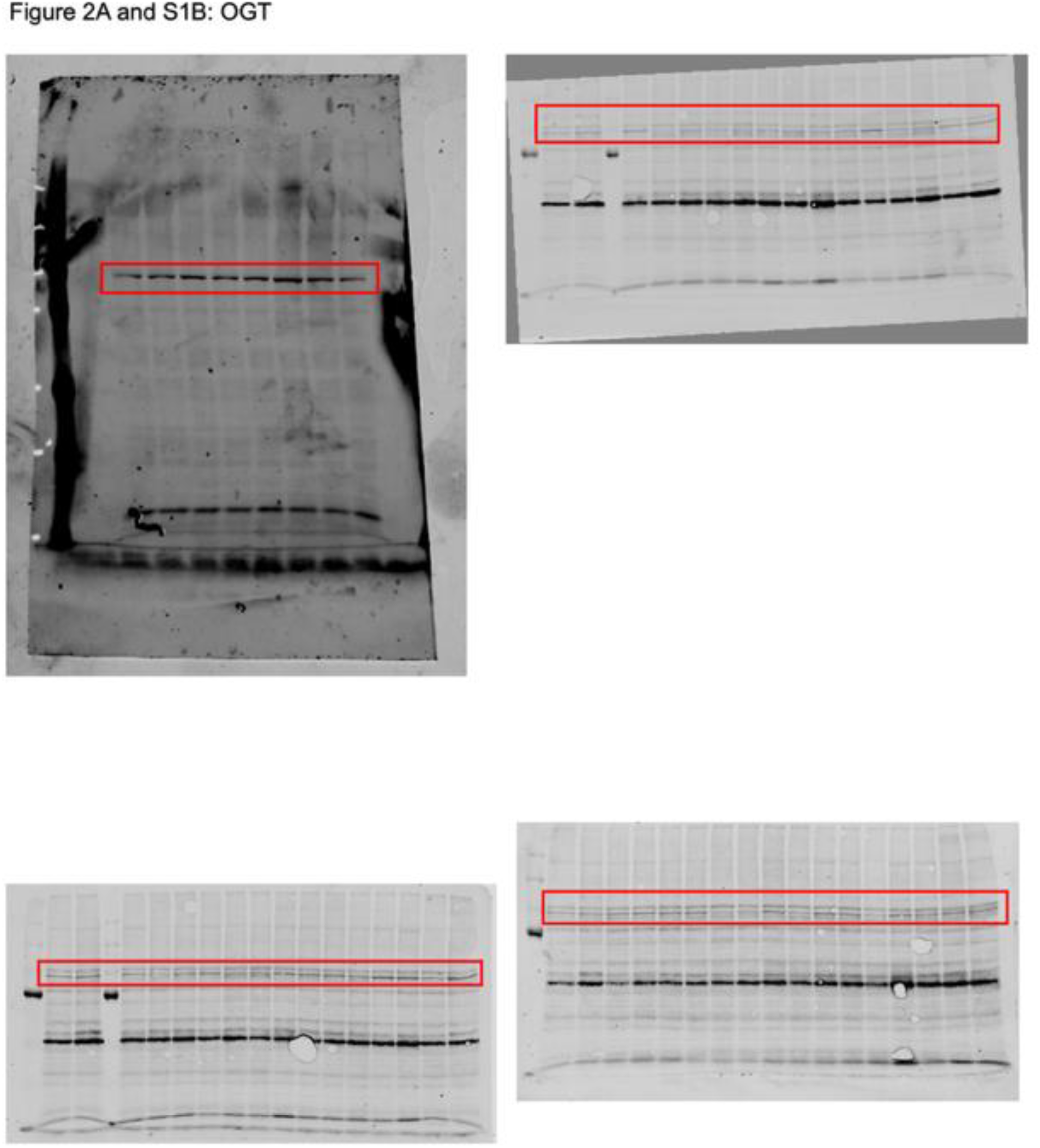

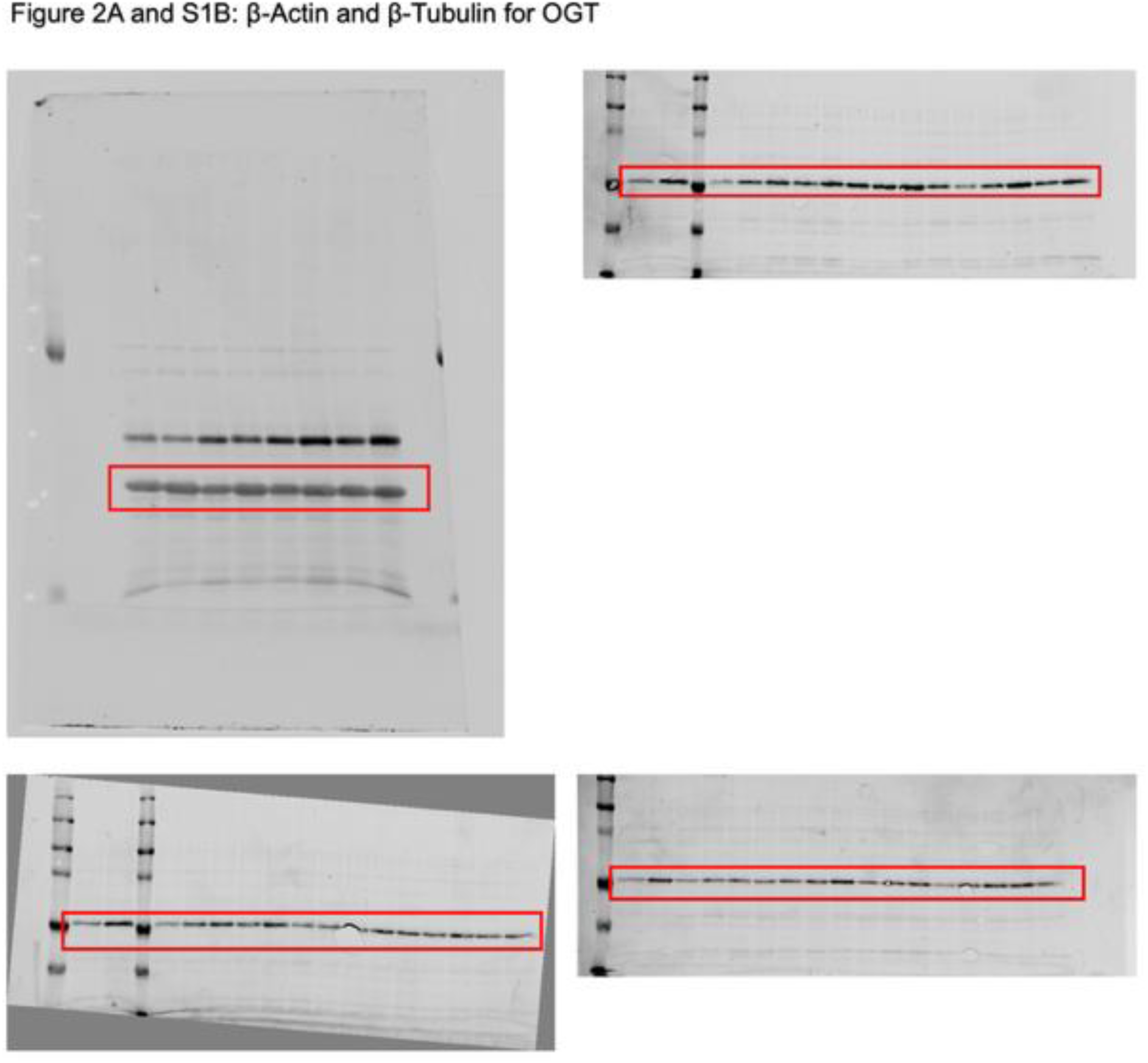

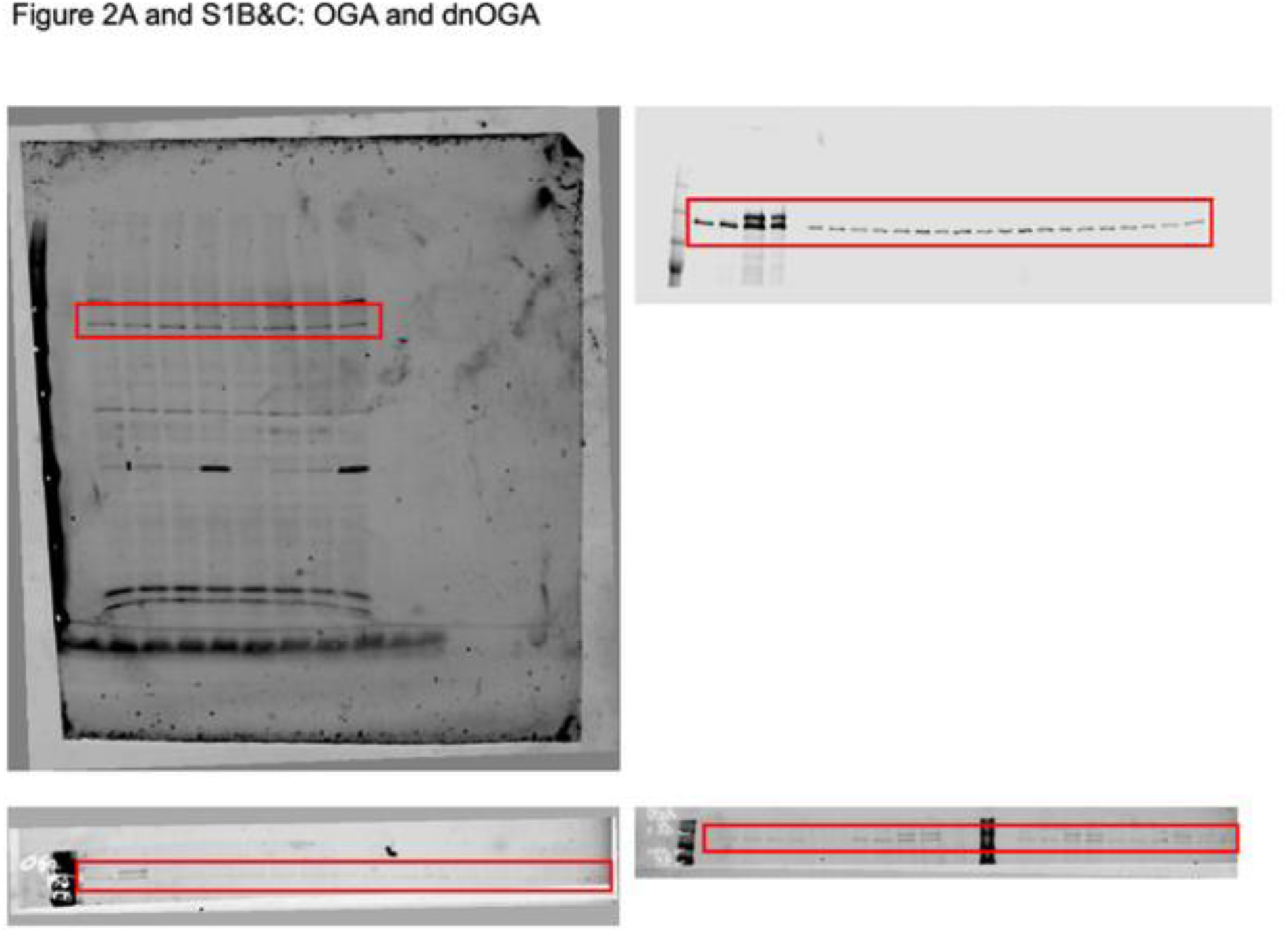

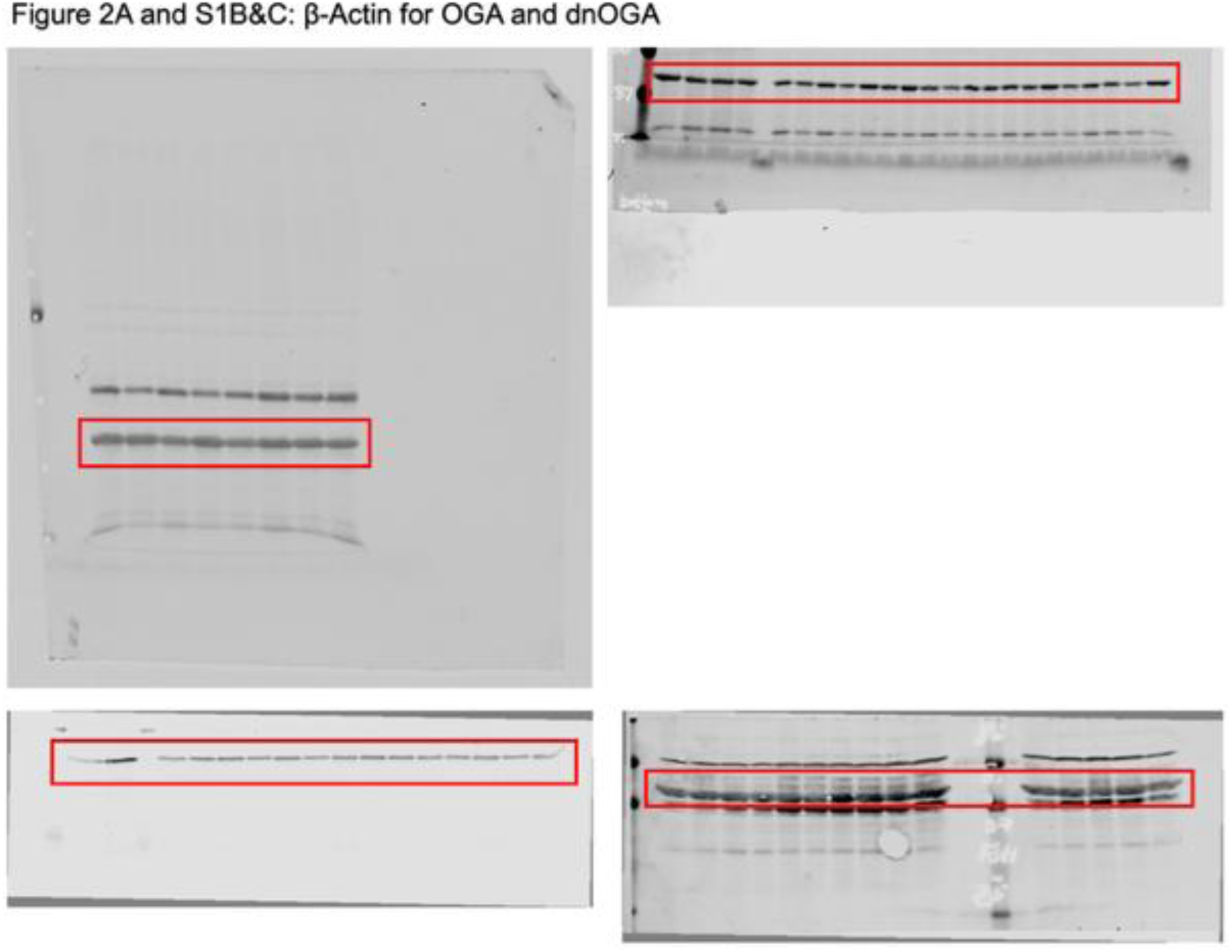

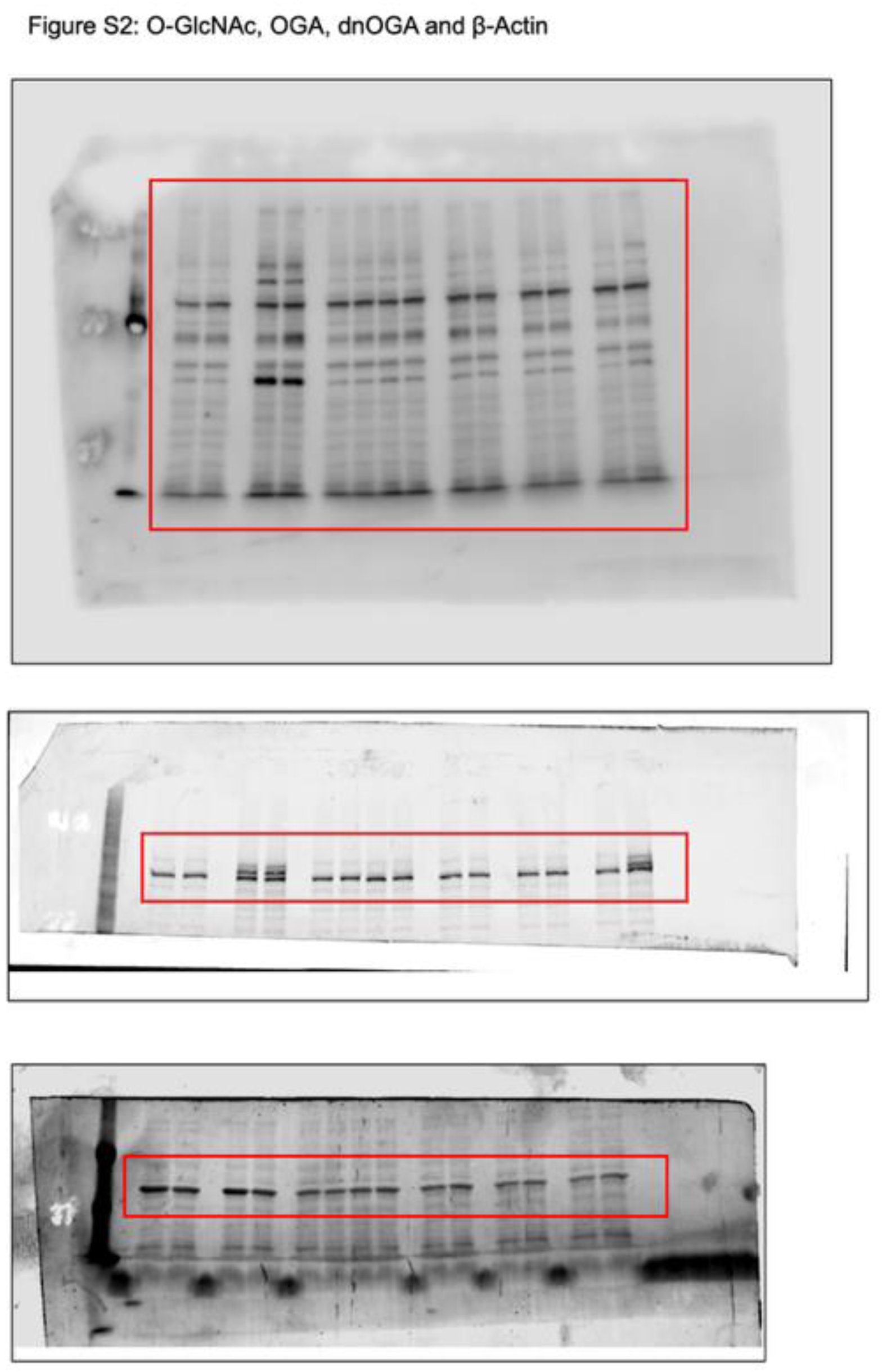

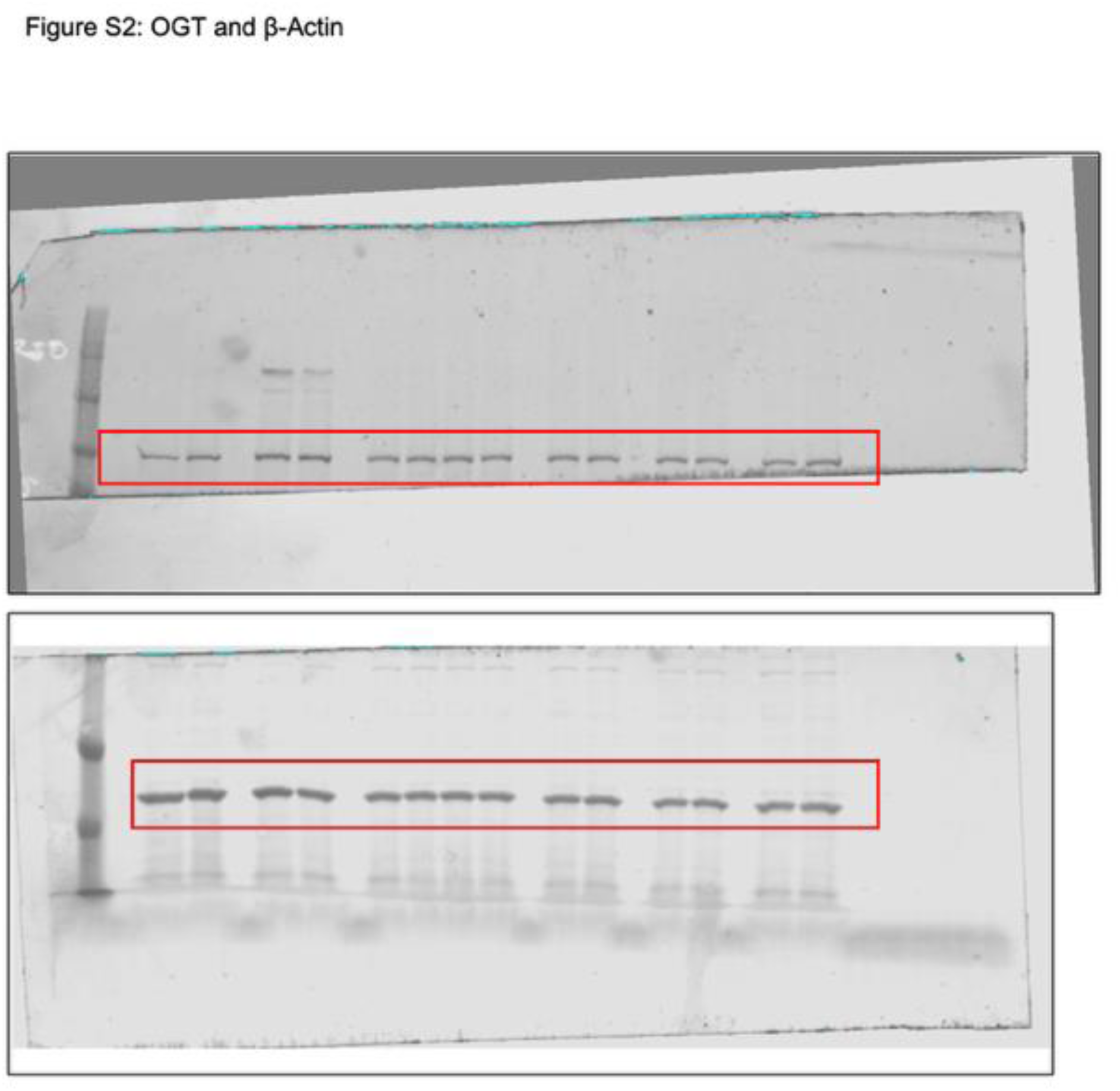

